# PTEN Deletion in the Adult Dentate Gyrus Induces Epilepsy

**DOI:** 10.1101/2024.08.07.606938

**Authors:** Jennifer M. Yonan, Kevin D. Chen, Tallie Z. Baram, Oswald Steward

**Author notes:** Address correspondence to: Oswald Steward, Ph.D., 1105 GNRF, 837 Health Sciences Dr., University of California at Irvine, Irvine, CA 92697.

## Abstract

Embryonic and early postnatal promotor-driven deletion of the phosphatase and tensin homolog (PTEN) gene results in neuronal hypertrophy, hyperexcitable circuitry and development of spontaneous seizures in adulthood. We previously documented that focal, vector-mediated PTEN deletion in mature granule cells of adult dentate gyrus triggers dramatic growth of cell bodies, dendrites, and axons, similar to that seen with early postnatal PTEN deletion. Here, we assess the functional consequences of focal, adult PTEN deletion, focusing on its pro-epileptogenic potential. PTEN deletion was accomplished by injecting AAV-Cre either bilaterally or unilaterally into the dentate gyrus of double transgenic PTEN-floxed, ROSA-reporter mice. Hippocampal recording electrodes were implanted for continuous digital EEG with concurrent video recordings in the home cage. Electrographic seizures and epileptiform spikes were assessed manually by two investigators, and corelated with concurrent videos. Spontaneous electrographic and behavioral seizures appeared after focal PTEN deletion in adult dentate granule cells, commencing around 2 months post-AAV-Cre injection. Seizures occurred in the majority of mice with unilateral or bilateral PTEN deletion and led to death in several cases. PTEN-deletion provoked epilepsy was not associated with apparent hippocampal neuron death; supra-granular mossy fiber sprouting was observed in a few mice. In summary, focal, unilateral deletion of PTEN in the adult dentate gyrus suffices to provoke time-dependent emergence of a hyperexcitable circuit generating hippocampus-origin, generalizing spontaneous seizures, providing a novel model for studies of adult-onset epileptogenesis.

## 1. Introduction

Epilepsy is a complex, multifactorial entity, with both genetic and environmental origins. Among genetic risk factors for epilepsy, dysregulation of the mechanistic target of rapamycin (mTOR) pathway, as takes place in tuberous sclerosis, is well-established. However, the mTOR pathway consists of numerous key enzymes with protean cellular functions. Thus, it is unclear which deficits in the distinct components of the pathway lead to epilepsy. Mutations of the phosphatase and tensin homolog (PTEN) gene, an important upstream negative regulator of the mTOR pathway, are one candidate, motivating studies of consequences of targeted mutations of PTEN in murine models.

The general strategy for studies of PTEN has been to use promoter-driven Cre expression in PTEN-floxed mice to genetically delete PTEN in particular populations of neurons and/or glia in early development. For example, when deletion is driven by the GFAP promoter, PTEN is deleted in astrocytes and neurons in widespread brain regions during embryonic development. In this situation, there is dramatic brain and neuronal hypertrophy accompanied by progressive development of spontaneous seizures and early mortality (Backman et al., 2001; Fraser et al., 2008; Fraser et al., 2004; Kwon et al., 2003; Kwon et al., 2001). Subsequent studies have assessed the consequences of deleting PTEN in the early postnatal period selectively in newborn granule cells of the dentate gyrus by driving Cre expression either under the Gli1 promoter or via intradentate retroviral injection, for example. These studies also documented dramatic neuronal hypertrophy in the dentate gyrus, including enlarged granule cell bodies, increased dendrite complexity, and aberrant expansion of mossy fiber axon connectivity (Arafa et al., 2019; LaSarge et al., 2015; Pun et al., 2012; Williams et al., 2015). Physiological consequences of these morphological changes have been reported to include the development of spontaneous seizures and the formation of hyperexcitable, pro-epileptic hippocampal circuits (LaSarge et al., 2016; Pun et al., 2012; Santos et al., 2017; Williams et al., 2015).

In contrast to previous studies of deleting PTEN during development, our lab has focused on consequences of deleting PTEN in fully-mature neurons in adult rodents. Of note, we discovered that both AAV-Cre mediated PTEN deletion in PTEN-floxed adult mice (Gallent & Steward, 2018) and AAV-shRNA mediated PTEN knockdown in adult rats (Steward et al., 2019) results in re-initiation of a growth phenotype in mature cortical neurons involving increases in cell body size and dendritic arborization. More recently, we have expanded our studies to consequences of focal deletion of PTEN in the mature dentate gyrus of adult mice. Injections of AAV-Cre into the dentate gyrus of adult PTEN-floxed mice resulted in focal PTEN deletion, triggering growth of granule cell bodies, elongation of dendrites, robust formation of additional spines (and presumably new synapses) on elongated dendritic segments, and expansion of mossy fiber terminal fields in target areas (Yonan & Steward, 2023). In this collection of studies, casual observations did not reveal spontaneous behavioral seizures out to 6 months following PTEN deletion. However, two mice died during the night after exhibiting no signs of ill health and these sudden deaths might have occurred during a spontaneous seizure. Despite the fact that no seizures were observed in these mice, abnormal physiological activity including electrographic seizures cannot be excluded without continuous recording and monitoring.

The present study assessed the functional consequences of focal PTEN deletion in the adult dentate gyrus using continuous video-EEG focusing specifically on the development of spontaneous seizures (epileptogenesis). We find that both bilateral and unilateral vector-mediated PTEN deletion in adult PTEN-floxed, ROSA-reporter mice lead to the development of spontaneous seizures beginning around 2 months after AAV-Cre injection. Of note, there were three instances of sudden death during a prolonged seizure (SUDEP). Unlike excitotoxin models of epilepsy, histological assessments revealed no apparent loss of hippocampal neurons in CA3. Thus, focal PTEN deletion provides a novel, toxin/convulsant-free model of adult-onset temporal lobe epilepsy (TLE) in which the pathophysiology is initiated by a localized focus within the hippocampus.

## 2. Materials and methods

### 2.1 Experimental mice

Experiments involved two transgenic strains of mice developed in our local breeding colony. The first strain was generated by crossing PTEN-floxed mice carrying lox-p flanked exon 5 of the PTEN gene (RRID: IMSR_JAX:004597) with ROSA26tdTomato (Rosa^tdTomato^) reporter mice having a lox-P flanked STOP cassette in the Rosa locus upstream of a tandem dimer tomato (tdT) fluorescent protein sequence (RRID: IMSR_JAX:007905). This double transgenic strain is designated PTEN^f/f^/Rosa^tdTomato^. All studies involved mice that were homozygous at the transgenic loci. One other line was created by crossing this strain with Thy1-eYFP mice, originally purchased from the Jackson Labs, to generate mice that were homozygous at the PTEN^f/f^/Rosa^tdTomato^ loci and hemizygous for Thy1-eYFP. For simplicity, these mice will be referred to as PTEN/tdT mice, regardless of their eYFP expression. All the strains used in these studies were generated in our lab, and therefore have different genetic backgrounds than the original mice from Jackson Labs.

### 2.2 Surgical procedure, AAV-Cre injections into the dentate gyrus

All experimental procedures were approved by the Institutional Animal Care and Use Committee (IACUC) at the University of California, Irvine. Studies involved adult mice (at least 2 months of age at the time of AAV-Cre injection). Briefly, mice were anesthetized with Isoflurane (2-2.5%), placed in a stereotaxic device, and small cranial window was created above the injection site. Using a 10μl Hamilton syringe with a pulled glass pipette tip, either a single unilateral or bilateral injection of AAV2-Cre (Vector Bio Labs, 7011) or AAV2-GFP (Vector Bio Labs, 7004) was made at +/−1.3mm lateral and +2.2mm anterior to lambda at a depth of −1.6mm from the cortical surface. Each injection was 0.6μl in volume (1×10^12^ genome copies (GC)/ml in 1x phosphate-buffered saline (1xPBS, 20mM, pH 7.4) with 5% glycerol) and was performed over 4 minutes. The pipette was left in place for an additional 2 minutes before removal. Following completion of the surgery, mice were allowed to recover for 48 hours in a cage on a 37°C heating pad and then were returned to standard housing conditions. Surgeries were performed in four separate iterations, termed Cohorts, three injected with AAV-Cre and a fourth injected with AAV-GFP (Table 1).

**Table 1:** Animal numbers, attrition, and exclusions.

| Cohort | Vector | AAV Group | Died post AAV Sx | Died post EEG Sx | Excluded post death/perfusion | Final numbers |
| --- | --- | --- | --- | --- | --- | --- |
| 1 | AAV-Cre | Bilateral, n=11 | n=1 | n=2 | n=1, large brain lesion<br>n=2, ended recording early | n=5 |
| 2 | AAV-Cre | Unilateral, n=6 | n=0 | n=2 | n=1, incorrect EEG probe | n=3 |
| 3 | AAV-Cre | Unilateral, n=7 | n=0 | n=0 | n=0 | n=7 |
| 4 | AAV-GFP | Unilateral, n=3<br>Bilateral, n=3 | n=0<br>n=0 | n=0<br>n=0 | n=1, AAV missed<br>n=1, AAV missed | n=4 |

### 2.3 Surgical procedure: Bilateral and unilateral intrahippocampal EEG probe placement

At 4-6 weeks post AAV-Cre injection, mice underwent a second procedure for placement of either bilateral or unilateral intrahippocampal electrodes, as described previously (Chen et al., 2021; Garcia-Curran et al., 2019). Mice were anesthetized with Isoflurane (1.5-2.5%) and placed in a stereotaxic device. A scalp incision was made to expose the skull and the skull was cleaned and dried with a 30% hydrogen peroxide solution. Two burr holes were placed in the skull above the hippocampus in each hemisphere at +/−1.6mm lateral and −1.9 posterior to bregma. Twisted wire electrodes were placed at a depth of −0 to −2.2mm below the brain surface (P Technologies, E363/2-2TW/SPC). For mice with a unilateral EEG probe, the electrode was placed into the right hippocampus, contralateral to AAV-Cre injection. Another 2 burr holes were created above on the left side of the cerebellum and right frontal cortex for placement of ground screw electrodes (P technologies, E363/20). Each electrode was then placed into a 6-channel pedestal (P Technologies, MS363). Electrodes, screws, pedestals, and wiring were held in place using a combination of cyanoacrylate and dental cement for creation of a head cap. Following completion of the surgery, mice were allowed to recover in a cage on a 37°C heating pad and then were returned to standard housing conditions until transferred to the continuous recording chambers. Information regarding animal numbers, attrition and exclusions are listed in Table 1.

### 2.4 Continuous video EEG recordings

At 6-8 weeks post AAV-Cre injection (1-2 weeks following electrode placement), mice were transferred into plexiglass cages and attached to a commutator to allow for free movement during video-EEG recording. Continuous recordings and videos were collected using the Lab Chart EEG analysis software (versions 7 and 8, AD Instruments), as described previously (Chen et al., 2021; Dube et al., 2010). Bilateral recordings were made in mice that received bilateral electrodes. In mice that received unilateral injections and unilateral electrodes, recordings were made on the side contralateral to the injection. Information on recording durations for each cohort of mice is listed in Tables 2-5.

**Table 2:** Cohort 1, Bilateral PTEN-deleted mice with bilateral electrode placement used for EEG recordings.

| Animal | Strain | AAV-Cre location | EEG location | DPIs recorded | DPI first seizure | Total # seizures | DPI died/sacrificed |
| --- | --- | --- | --- | --- | --- | --- | --- |
| 307-8 | PTEN/tdT | Bilateral ^ | Bilateral | 43-113 | 65 | 42 | 113 |
| 307-9 | PTEN/tdT | Bilateral + | Bilateral | 43-113 | 79 | 27 | 113 |
| 329-20 | PTEN/tdT | Bilateral ^ | Bilateral | 42-187 | 87 | 33 | 190 |
| 329-21 | PTEN/tdT | Bilateral ^+ | Bilateral | 42-147 | 130 | 12 | S.D. 147 |
| 329-22 | PTEN/tdT | Bilateral ^ | Bilateral | 42-169 | n/a | 0 | 201 |

^ Bilateral injection with mild/moderate transduction of one/both CA1
+ Bilateral injection with one injection mistargeted to CA1
S.D.= seizure-related death

### 2.5 EEG analysis and scoring

Analyses of EEG recordings were accomplished by investigators blind to animal group. A subset of the EEGs were independently analyzed by two investigators, with excellent concordance. Recordings were scanned for seizures, defined as events lasting more than 6 seconds and consisting of EEG polyspikes or sharp-waves (amplitude > 2-fold background) (Chen et al., 2021; Dube et al., 2006; Pitkanen et al., 2002). In addition, the progression of the amplitude and the frequencies of the discharges throughout a given seizure were analyzed, because typical seizures are characterized by increasing amplitude and slowing frequency as the seizure progresses. Finally, seizures required a period of post-ictal depression characterized by a dramatic decrease in EEG amplitude. In mice with bilateral recording electrodes, both ipsilateral and contralateral EEG recordings were scored. In addition to electrographic analyses, videos accompanying seizures were analyzed as described (Dube et al., 2006; Dube et al., 2010). Briefly, we evaluated typical behaviors associated with limbic-onset seizures, including sudden cessation of activity, facial automatisms, head-bobbing, prolonged immobility with staring. These progressed to alternating or bilateral clonus, rearing and falling (Racine). Mice were considered epileptic if they had at least one documented spontaneous seizure as defined above. Data are represented as the percentage of mice within each cohort to develop seizures, cumulative number of seizures for each individual animal over time, the total number of seizures for each mouse, the latency to develop spontaneous seizures, and the average number of seizures per days recorded.

### 2.6 Tissue collection and histology

At designated endpoints (Tables 2-5), mice were euthanized via intraperitoneal injections of Fatal Plus® (390mg/ml pentobarbital sodium) and were intracardially perfused with 4% paraformaldehyde in phosphate buffer (PFA). Brains were dissected and post-fixed in 4% PFA for 48 hours, cryoprotected in 27% sucrose, and frozen in Tissue-Tek O.C.T. compound. Brains were then cryosectioned at 30μm and stored in 1xPBS with 0.1% NaN_3_ until processed for immunohistochemistry. For mice that were retrieved after being found dead, brains were drop-fixed in 4% PFA and then prepared as above. The area of transduction was visualized by tdTomato expression in PTEN/tdT mice that received AAV-Cre and GFP expression in control mice that received AAV-GFP.

Sets of sections at 360μm intervals were processed for different histological markers. For immunohistochemistry (IHC), sections were washed in tris-buffered saline (1xTBS, 100mM Tris, pH 7.4 and 150mM NaCl) then quenched for endogenous peroxidase activity by incubation in 3% H_2_O_2_ for 15 minutes. Sections were then washed in 1xTBS and blocked in blocking buffer (1xTBS, 0.3% Triton X-100, 5% normal donkey serum (NDS)) for 2 hours at room temperature. Sections were then incubated overnight at room temperature in buffer containing primary antibodies for rabbit anti-PTEN (1:250, Cell Signaling Technology 9188, RRID: AB_2253290), rabbit anti-pS6 Ser235/236 (1:250, Cell Signaling Technology 4858, RRID: AB_916156), rabbit anti-Znt3 (1:1500, Millipore ABN994), rabbit anti-cFos (1:1000, Millipore ABE457, RRID: AB_2631318), and mouse anti-GAD67 (1:1000, Millipore MAB5406, RRID: AB_2278725). Sections were then washed in 1xTBS, followed by a 2-hour incubation in buffer containing biotinylated donkey anti-rabbit IgG (1:250, Jackson ImmunoResearch, 711-065-152), then washed again. Visualization was accomplished through incubation in avidin-biotin complex (ABC) reagent (Vectastain Elite kit, catalog #PK-6100; Vector Laboratories) and catalyzed reporter deposition (CARD) amplification with Tyramide-FITC or Tyramide-AMCA. Sections stained for rabbit anti-GFP (1:1500, Novus NB600-308, RRID:10003058) were visualized using Alexa fluor-488 (1:250, Invitrogen A21206). Sections stained for GAD67 were visualized by DAB (Vector Laboratories, SK-4100), mounted on slides, dehydrated through graded ethanol, cleared in xylenes, and coverslipped with DPX). All fluorescently labeled sections were then mounted on 0.5% gelatin coated slides and counterstained with Hoechst (1μg/mL).

### 2.7 Extent of transduction of the dentate gyrus

To determine the percent area of transduction through the dentate gyrus, sections spaced 360μm apart were assessed for percentage of the granule cell layer occupied by tdTomato-positive granule cells as in our previous study (Yonan & Steward, 2023). Briefly, images of the dentate gyrus were taken with a 10x objective using an Olympus AX80 microscope. Images were imported into NIH ImageJ FIJI, a border was drawn around the Hoechst-positive granule cell layer, the drawn contour was transferred to the tdT-labeled image, and the tdT-positive area within the contour was determined. The percent of the granule cell layer that was tdT-positive throughout the rostro-caudal series of sections was calculated in each mouse. Data are plotted as percent transduction through the length of the dentate gyrus at 360μm intervals and percentage transduction of the entire dentate gyrus for each mouse.

The extent of PTEN deletion within each mouse was qualitatively assessed for transduction of the dentate gyrus and CA1 hippocampal subregion. These include mild/moderate transduction of the CA1 in one or both hippocampi in addition to the dentate gyrus or mistargeting of an injection in which the CA1 was transduced with minimal transduction of the dentate gyrus. Each of these distinctions is noted alongside each mouse in corresponding tables for each Cohort (Tables 2-5). Transduction of the CA1 that was isolated to the needle tract alone was classified as an injection of only the dentate gyrus.

### 2.8 Granule cell size measurements

Cell body sizes were measured in regions of dense transduction near the injection epicenter. One set of sections for each mouse was stained with Cresyl violet and a separate set of sections was assessed for the region of PTEN deletion through immunohistochemistry and visualization of tdTomato, as described above. A single section at the core of PTEN deletion and transduction was taken for granule cell size measurements. Z-stack images of Cresyl violet stained sections were taken using a 44.4x objective with a 1μm step size using the Keyence BZ-X800 microscope and imported into ImageJ FIJI. Sampling was done by measuring 30 cells within the granule cell layer within a 100 x 200 μm region of interest in the ipsilateral dentate gyrus, and the homologous regions in the contralateral granule cell layer. The mean cross-sectional area for PTEN deleted and PTEN expressing granule cells was determined by first averaging cell sizes from individual mice, then averaging for each hemisphere where n = 6 mice.

Measurements of the thickness of the molecular layer were taken at the core of transduction using images taken with a 10x objective on an Olympus AX80 microscope as a representation of maximal dendritic length (granule cell dendrites extend to the border of the molecular layer). Measures of molecular layer width from ipsilateral and contralateral sides were then compared.

### 2.9 Assessment of mossy fiber projections

Sections immunostained for the zinc transporter Znt3 were used to assess alterations in granule cell axonal projections (mossy fibers). Images of both the ipsilateral and contralateral dentate gyrus and CA3 were taken with a 10x objective using an Olympus AX80 microscope and imported into ImageJ FIJI. Measurements were taken of the thickness of the laminae containing mossy fibers as they exited the hilus on both ipsilateral and contralateral sides.

To detect supra-granular mossy fibers, Znt3 labeled sections at the core of transduction were used. Sections were scored as described by (Hunt et al., 2009): 0=little to no Znt3 labeling in granule cell layer; 1=mild Znt3 labeling in granule cell layer; 2=moderate staining in the granule cell layer and punctuate staining in inner molecular layer; 3=dense Znt3 labeling in inner molecular layer.

### 2.10 Statistical methods

Analyses were conducted without explicit knowledge of experimental group when feasible (in most cases, the area of transduction was obvious due to growth of the dentate gyrus). Graphs were created and statistical analyses were performed using GraphPad Prism Software. One-way analysis of variance (ANOVA) was used to compare seizure outcomes across cohorts. Two-way ANOVA was used for comparison of neuronal morphological measures between contralateral and ipsilateral hemispheres in PTEN/tdT mice. Sidak’s multiple comparisons tests were used for comparisons between groups. Relationships between neuronal outcome measures, percent PTEN deletion, or seizure number were assessed by linear regression.

## 3. RESULTS

### 3.1 Effective PTEN deletion in PTEN/tdT mice following unilateral and bilateral AAV-Cre injections into the dentate gyrus

Injections of AAV-Cre into the dentate gyrus of PTEN/tdT mice at 2 months of age resulted in robust transduction of mature granule cells surrounding the injection site. Figure 1 illustrates representative images of mice with bilateral and unilateral AAV-Cre injections collected between 2 and 4 months after injection. Figures 1A and 1B depict the regions of granule cell transduction following bilateral AAV-Cre injection based on tdTomato expression. In both dentate gyri, PTEN immunostaining was absent in the area of tdT expression (Fig. 1C, D). Immunostaining for the phosphorylated form of ribosomal protein S6 (a downstream marker of mTOR activation) revealed robust activation of S6 phosphorylation in the regions of PTEN deletion (Fig. 1E, F).

**Figure 1.**
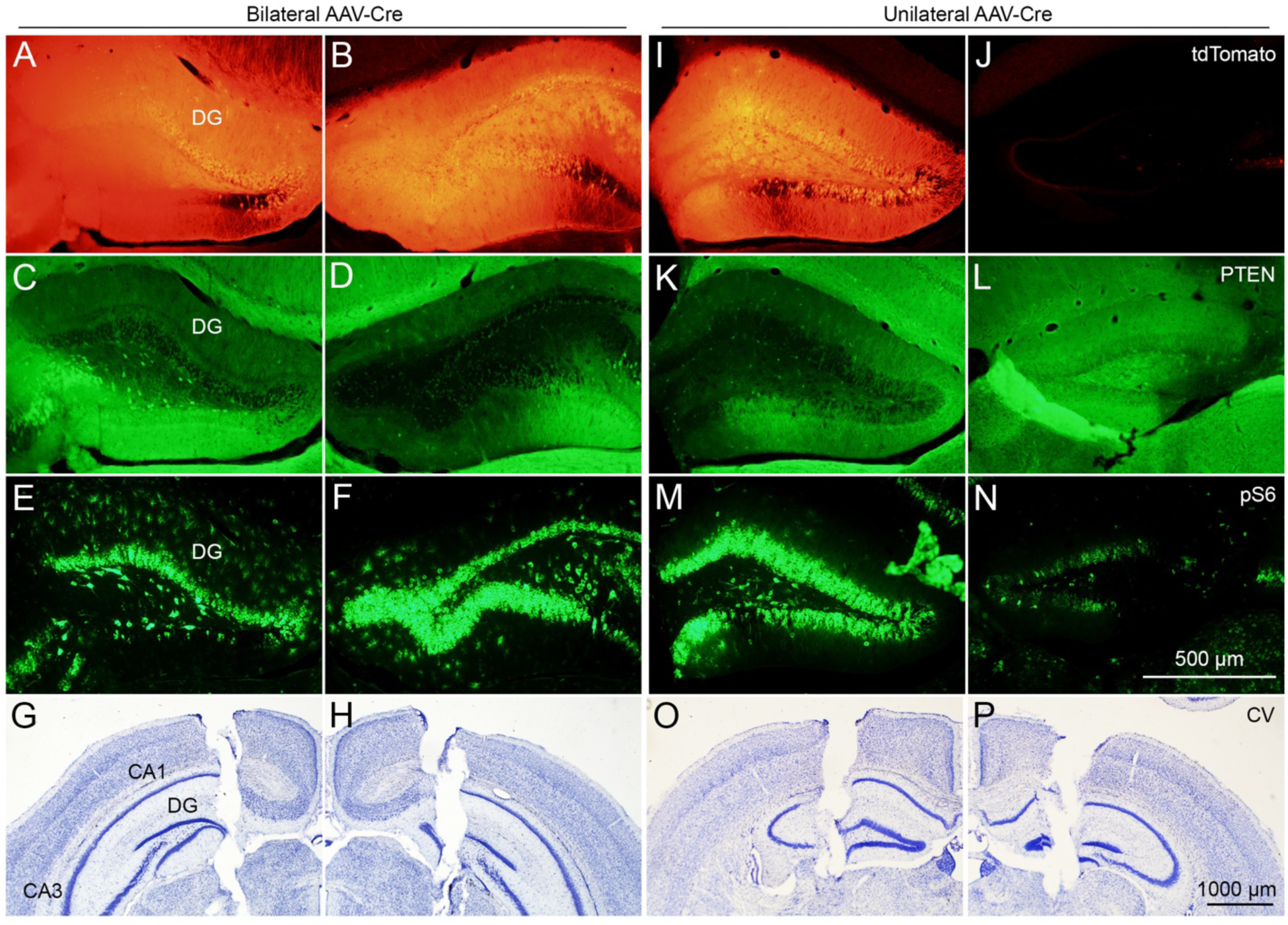
Effective PTEN deletion and mTOR activation in PTEN/tdT mice following bilateral and unilateral AAV-Cre injections into the dentate gyrus. A-B) tdTomato expression in transduced dentate granule cells following bilateral AAV-Cre injection in a PTEN/tdT mouse. C-D) Immunostaining for PTEN reveals deletion in tdT positive granule cells. E-F) Immunostaining for phospho-S6 indicates activation of mTOR in PTEN deleted granule cells. G-H) Cresyl violet stained sections reveal bilateral electrode placement into the hippocampus. I-J) tdT expression in the transduced dentate gyrus (left) and lack of tdT expression in the non-injected dentate gyrus (right) of a PTEN/tdT mouse following unilateral AAV-Cre injection. K-L) PTEN deletion in the ipsilateral dentate gyrus. Note, the preservation of PTEN expression in the contralateral dentate gyrus. M-N) pS6 immunoreactivity in the PTEN deleted and PTEN expressing dentate gyrus of the same mouse. O-P) Location of bilateral, intrahippocampal electrodes in the same mouse.

Figures 1I and 1J depict the regions of transduction following unilateral injection of AAV-Cre into the left dentate gyrus (compare tdT expression in Fig. 1I with the lack of tdT expression in the contralateral, non-injected dentate gyrus in Fig. 1J). PTEN deletion was confirmed by lack of immunostaining in transduced, tdT-positive granule cells (Fig. 1K), but remained intact in granule cells of the contralateral, non-injected dentate gyrus (Fig. 1L). Increases in immunostaining for pS6 were evident on the side of PTEN deletion (Fig. 1M) in comparison to the contralateral dentate gyrus (Fig. 1N).

### 3.2 Spontaneous seizures develop following bilateral PTEN deletion

#### Cohort 1, Bilateral PTEN deletion with bilateral EEG recording

Promoter-driven PTEN deletion in the developing brain results in PTEN deletion throughout the dentate gyri of both hemispheres (Kwon et al., 2001; Matsushita et al., 2016; Pun et al., 2012). To determine if bilateral but focal PTEN deletion in mature granule cells of the adult dentate gyrus results in pro-epileptic alterations to circuit function, recording electrodes were implanted bilaterally into the hippocampi 4-6 weeks after bilateral injections of AAV-Cre. Figure 1G, H illustrate examples of recording electrode tracks.

#### Animal attrition and exclusions

In Cohort 1 (Table 1), one mouse died prior to the EEG electrode placement surgery; tissue for this mouse was unavailable for postmortem analysis because the mouse was found dead by vivarium staff, and the carcass was disposed of. Two of 11 mice (18%) died at 3 and 4 days after electrode probe placement but prior to the initiation of EEG recordings. For 1 mouse, tissue was unavailable for analysis because the mouse was found dead, and the carcass was disposed of. For the remaining mouse, the brain was drop-fixed and was prepared for histology, which revealed appropriate transduction of granule cells in both hemispheres, with no transduction of other hippocampal subregions. The cause of death in these mice is unknown given that death occurred prior to initiation of EEG/video recordings. Another 1 of 11 mice (9%) exhibited seizures during the recording period but was excluded following tissue analysis due to a large brain lesion of unknown origin that may have influenced seizure onset and incidence.

#### Seizure onset and incidence

Four of five mice in which EEG was recorded (80%) developed spontaneous electrographic and behavioral seizures (Fig, 7A). The first seizures were observed at an average of 90.25 +/− 28.02 days (median = 83 days) post AAV-Cre injection (Fig. 7B). These mice experienced an average of 28.50 +/− 12.61 total seizures (median = 30 seizures) over the recording period (Fig. 2A, 7C, Table 2) with an average of 0.91 +/− 0.098 seizures per day (median = 0.94 seizures per day, Fig. 7D). Mice with bilateral PTEN deletion exhibited seizures that originated in either hippocampus (Fig. 2B, C). The frequency of the seizures did not increase over time (Fig. 2A) and seizures occurred in clusters. This is in contrast to reports with developmental PTEN deletion in which seizures increase in frequency and severity over time (Kwon et al., 2001). In this cohort, one mouse with bilateral PTEN deletion died at 147 days post AAV-Cre injection during a prolonged focal-onset, generalized seizure, after exhibiting a total of 12 seizures during the recording period (Fig. 2A, animal 329-21).

**Figure 2.**
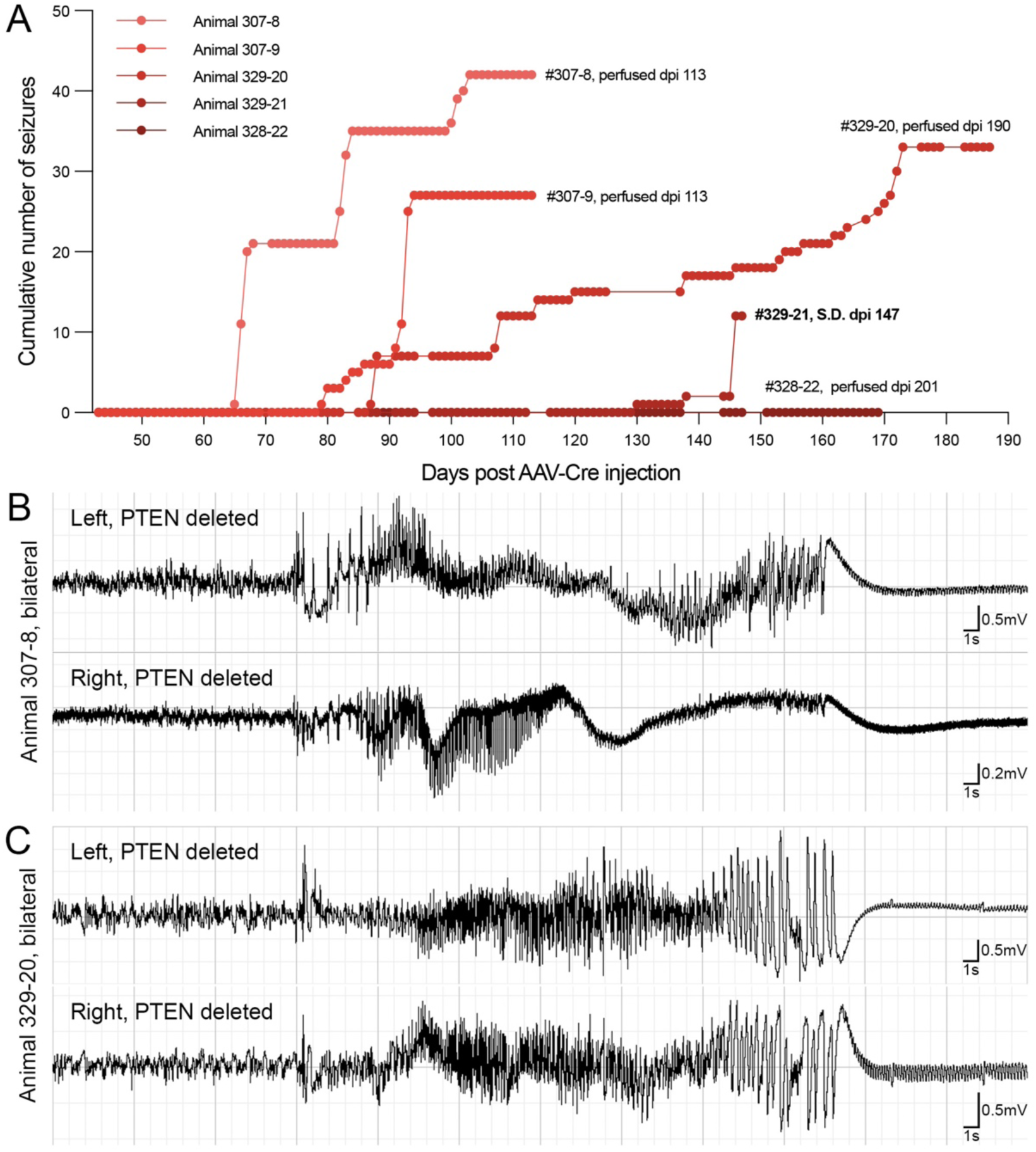
EEG recordings in PTEN/tdT mice (Cohort 1) following bilateral AAV-Cre injections into the dentate gyrus. A) Cumulative number of seizures for each mouse over time following bilateral PTEN deletion. B-C) Representative EEG recordings from two mice with bilateral PTEN deletion in the dentate gyrus. Note the typical progression of seizure activity in both hippocampi followed by a period of post ictal depression.

Interestingly, the majority of mice that were recorded for longer than 2 months after PTEN deletion developed seizures, suggesting an approximately 2-3 month latency period for development of epilepsy. In Cohort 1, two mice were recorded for only two-months and were sacrificed early to verify electrode placement (Table 1). Whether these 2 mice would have developed seizures had they been recorded for longer periods of time is unknown, so these two mice are excluded from the calculation of seizure prevalence.

#### Patterns of AAV-Cre transduction in PTEN/tdT mice with bilateral PTEN deletion

There was some variability in transduction efficacy despite the use of consistent stereotaxic coordinates, injection parameters, and vector titers. Table 2 provides detailed descriptions of the pattern of transduction and PTEN deletion in each mouse, specifically the accuracy of transduction of the dentate gyrus and the transduction of other hippocampal subregions. Three mice with bilateral transduction also had mild transduction of the CA1 region in one or both hippocampi, one mouse had bilateral transduction in which targeting in the right hippocampus was predominately in the CA1 region, and one mouse with bilateral transduction displayed mild transduction of the left CA1 and targeting in the right hippocampus predominately of the CA1. Of note, this final animal is the one that died during a verified seizure. It remains to be determined how these variations and patterns of PTEN deletion impact circuit and network function.

### 3.3 Spontaneous seizures develop following unilateral PTEN deletion

#### Cohort 2: Unilateral PTEN deletion with bilateral EEG recording

Temporal lobe epilepsy in humans often develops from a unilateral seizure focus. To determine if unilateral PTEN deletion is sufficient for seizure development, we carried out a pilot study in which 6 mice received unilateral injections of AAV-Cre into the dentate gyrus followed by placement of bilateral intrahippocampal electrodes. Figures 1O and P illustrate electrode tracks in a mouse with unilateral PTEN deletion.

#### Patterns of transduction

In Cohort 2, three of 6 mice (50%) were excluded from analysis due to death (two mice at 1- and 47-days post EEG implantation surgery) or due to incorrect electrode placement (Table 1). Two of three mice included in analysis (Table 3), as well as mice with tissue available postmortem, all displayed appropriate and well targeted transduction of only the dentate gyrus. In these cases, PTEN expression was retained in both the CA1 and CA3. One of three mice included in analysis displayed some transduction of the CA1.

**Table 3:**
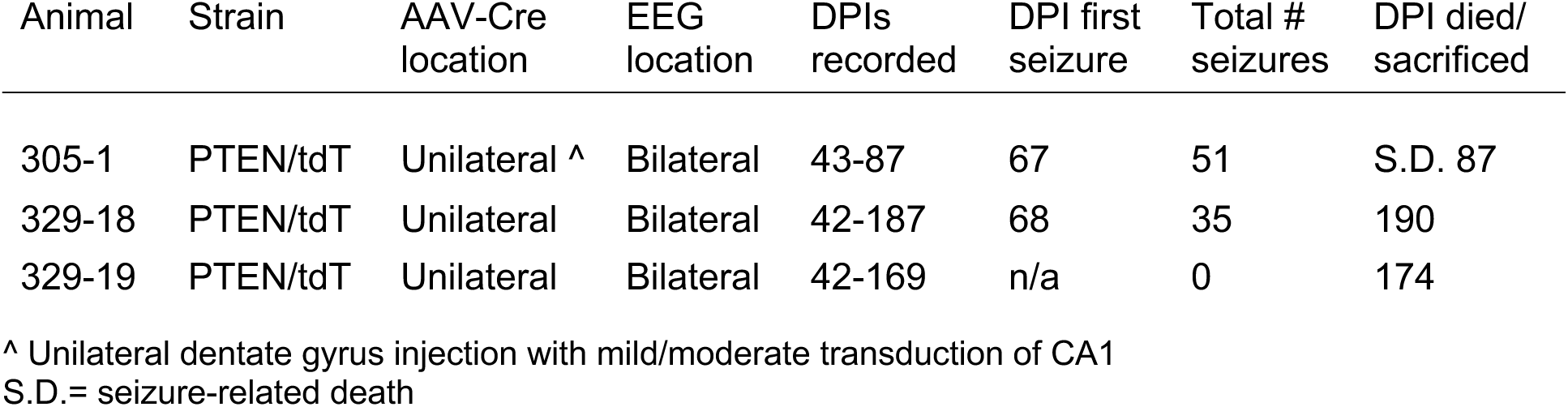
Cohort 2, Unilateral PTEN-deleted mice with bilateral electrode placement used for EEG recordings.

#### Seizure onset and frequency

In cohort 2, 2 of 3 mice (67%) developed spontaneous seizures (Fig. 7A) at an average of 67.50 +/− 0.71 days post AAV-Cre injection (median = 67.50 days), having an average of 43.00 +/− 11.31 seizures (median = 43 seizures) over the recording period, and an average of 1.60 +/− 1.75 seizures per day (median = 1.60 seizures per day) (Fig. 3A, Fig. 7B-D). In mice with unilateral PTEN deletion, seizures consistently originated in the PTEN deleted dentate gyrus and then propagated to the contralateral, PTEN-expressing dentate gyrus (Fig. 3B, C). As seen with bilateral PTEN deletion, seizures were clustered, did not increase in frequency over time, and began more than 2 months post PTEN deletion. In this cohort, animal 305-1 displayed the greatest number of total seizures over time, eventually dying suddenly at 87 days post injection during a prolonged seizure, having experienced a total of 51 seizures during the recording period.

**Figure 3.**
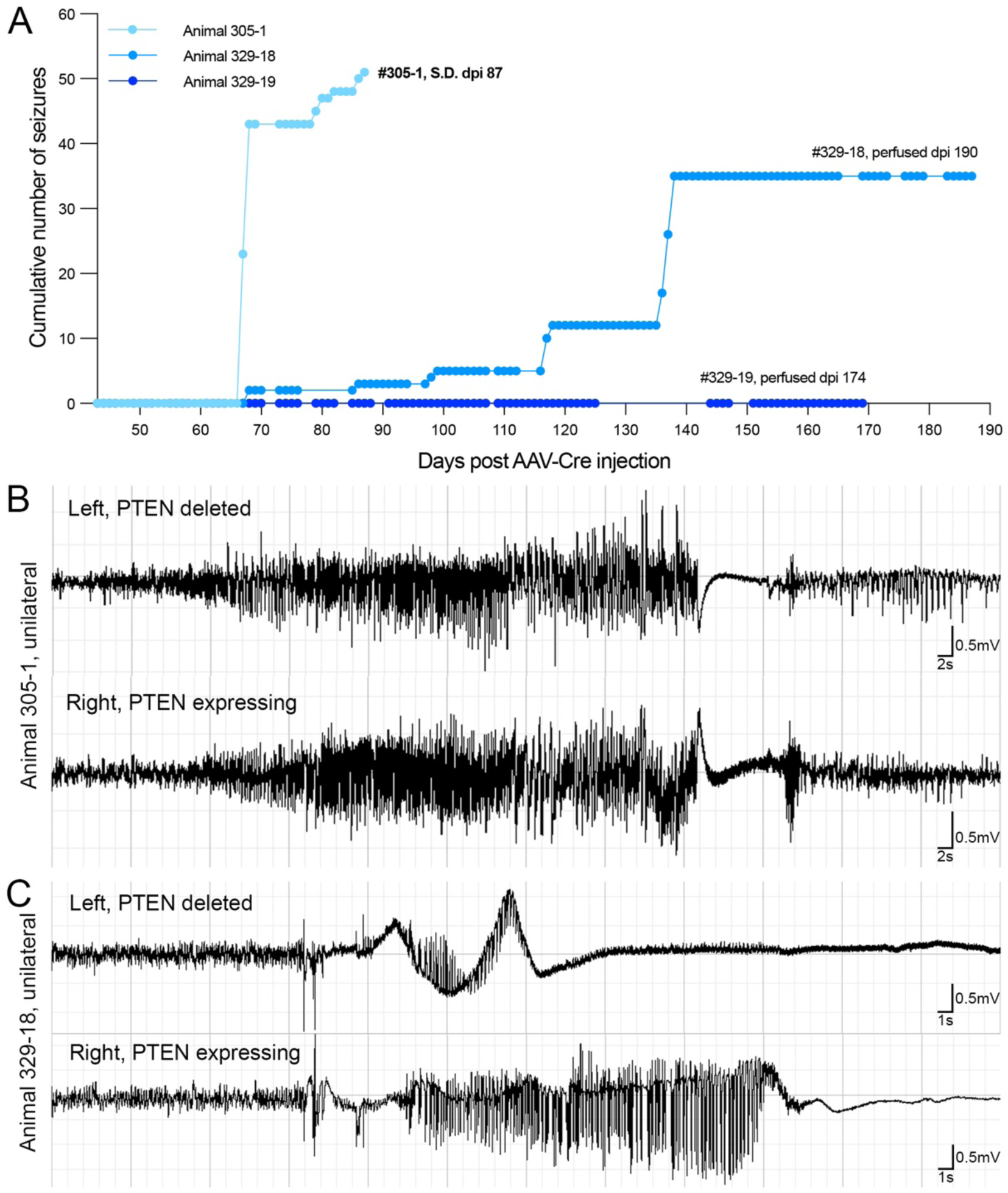
EEG recordings in PTEN/tdT mice (Cohort 2) following unilateral AAV-Cre injections into the dentate gyrus. A) Cumulative number of seizures for each mouse over time with unilateral PTEN deletion. B-C) Representative EEG recordings from two mice with unilateral PTEN deletion in the dentate gyrus. Note, seizure activity is initiated in the PTEN deleted hippocampus then propagates to the contralateral hippocampus.

### 3.4 Spontaneous seizures develop following unilateral PTEN deletion with contralateral EEG probe placement

#### Cohort 3, unilateral PTEN deletion with unilateral EEG recording

It was noteworthy that several mice in cohorts 1 and 2 died suddenly, and in 2 cases, death occurred during a prolonged seizure. In our previous studies of anatomical consequences of PTEN deletion in the adult dentate gyrus, in which mice survived up to 6 months after PTEN deletion (Yonan & Steward, 2023), behavioral seizures were not observed and there were only two instances of sudden death. Because we had not previously seen high mortality rates in mice with PTEN deletion but no implanted recording electrodes, we wondered whether PTEN deletion and electrode placement within the same regions could synergistically trigger more severe seizures resulting in increased incidence of sudden death.

To address this, PTEN/tdT mice (Cohort 3, Tables 1 and 4) received unilateral injections of AAV-Cre into the dentate gyrus followed by unilateral EEG electrode placement into the contralateral dentate gyrus (Fig. 4G, H). As above, AAV-Cre injection resulted in expression of tdTomato in transduced granule cells in the ipsilateral dentate gyrus (Fig. 4A vs. 4B), PTEN deletion in transduced cells alone (Fig. 4C vs. 4D), and activation of S6 phosphorylation in PTEN deleted granule cells (Fig. 4E vs. 4F). Notably, in Cohort 2, seizures recorded in mice with unilateral AAV-Cre injections always originated in the PTEN deleted dentate gyrus and consistently propagated to the contralateral hippocampus (Fig. 3B, C). Therefore, in this paradigm seizures recorded in the contralateral dentate gyrus are likely propagated from the PTEN-deleted dentate gyrus.

**Figure 4.**
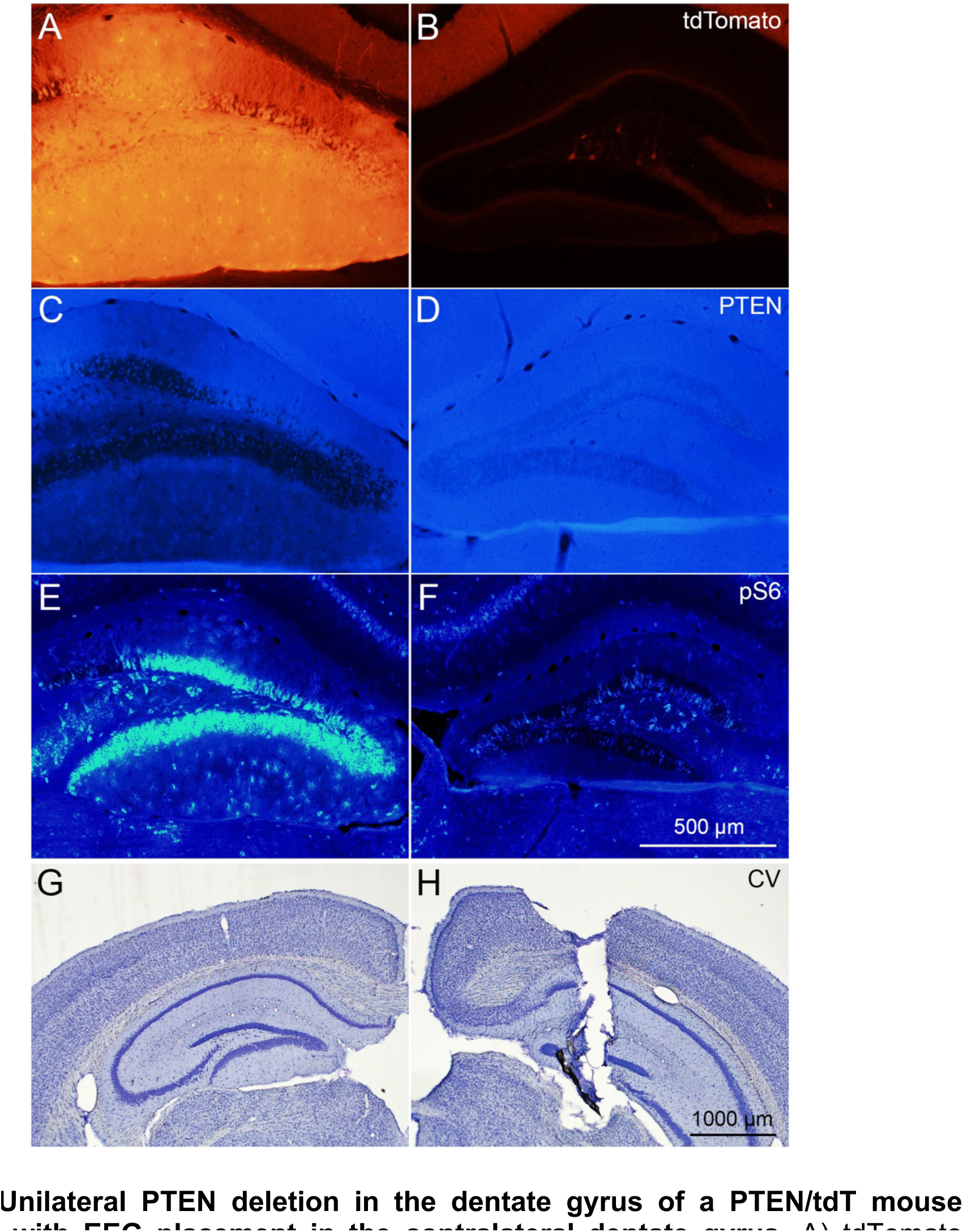
Unilateral PTEN deletion in the dentate gyrus of a PTEN/tdT mouse (Cohort 3) with EEG placement in the contralateral dentate gyrus. A) tdTomato expression in transduced granule cells of the ipsilateral dentate gyrus following unilateral AAV-Cre injection into a PTEN/tdT mouse. B) Lack of tdT expression in non-transduced cells in the contralateral dentate gyrus. C) PTEN deletion in tdT positive granule cells. D) Preservation of PTEN expression in the contralateral dentate gyrus. E) Increased phosphorylation of ribosomal protein S6 in PTEN deleted granule cells. F) pS6 immunoreactivity in the contralateral dentate gyrus. G) Cresyl violet stained section at the core of transduction and PTEN deletion. H) Cresyl violet stained section showing unilateral electrode placement into the contralateral hippocampus.

#### Seizure onset and incidence

Mice were recorded for 4 months post injection. Seven of seven mice recorded developed spontaneous seizures at varying frequency (Table 4, Fig. 5, Fig. 7A). The average time to first seizure in this cohort of mice was 69.29 +/− 5.68 days (median = 71 days) with an average of 28.43 +/− 24.64 total seizures (median = 23 seizures) over the recording period, and 1.35 +/− 2.09 seizures per day (median = 0.75 seizures per day). One mouse died suddenly at 77 days post AAV-Cre injection during a seizure, having had a total of 12 seizures (Fig. 5A, animal 69-29). These results strongly support the conclusion that unilateral PTEN deletion in the mature dentate gyrus is sufficient to result in the formation of a circuit that eventually leads to spontaneous electrographic and behavioral seizures. While not an entirely isolated circuit, this paradigm mostly eliminates the possibility of a two-hit model of seizure initiation.

**Figure 5.**
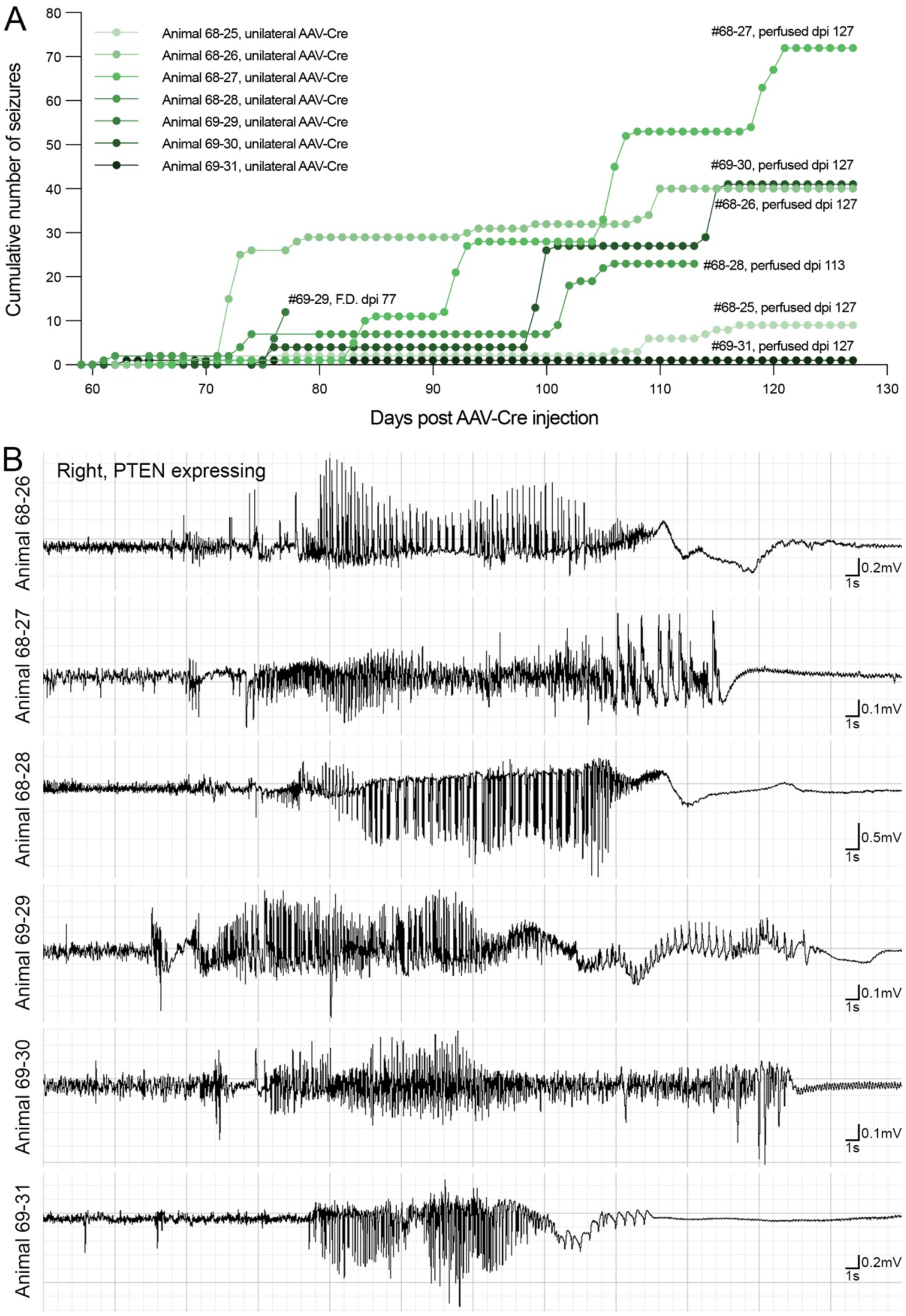
EEG recordings in PTEN/tdT mice (Cohort 3) following unilateral AAV-Cre injections into the dentate gyrus and placement of EEG electrode into contralateral hippocampus. A) Cumulative number of seizures recorded in the contralateral, PTEN-expressing dentate gyrus for each mouse over time. B) Representative EEG recordings from all mice with unilateral PTEN deletion in the dentate gyrus.

**Table 4:** Cohort 3, Unilateral PTEN-deleted mice with contralateral electrode placement used for EEG recordings.

| Animal | Strain | AAV-Cre location | EEG location | DPIs recorded | First Seizure | Total # seizures | DPI died/ sacrificed |
| --- | --- | --- | --- | --- | --- | --- | --- |
| 68-25 | PTEN/tdT/Thy1 | Unilateral | contralateral | 56-127 | 71 | 9 | 127 |
| 68-26 | PTEN/tdT/Thy1 | Unilateral | contralateral | 56-127 | 71 | 41 | 127 |
| 68-27 | PTEN/tdT/Thy1 | Unilateral | contralateral | 56-127 | 68 | 72 | 127 |
| 68-28 | PTEN/tdT/Thy1 | Unilateral | contralateral | 59-113 | 61 | 23 | 113 |
| 69-29 | PTEN/tdT/Thy1 | Unilateral | contralateral | 59-77 | 76 | 12 | S.D. 77 |
| 69-30 | PTEN/tdT/Thy1 | Unilateral ^ | contralateral | 59-127 | 75 | 41 | 127 |
| 69-31 | PTEN/tdT/Thy1 | Unilateral ^ | contralateral | 62-127 | 63 | 1 | 127 |

^ Unilateral dentate gyrus injection with mild/moderate transduction of CA1 S.D.= seizure-related death

### 3.5 Unilateral or bilateral AAV-GFP injections do not trigger seizures

Recently, a toxic effect of AAV virus injection on dentate gyrus granule cells has been reported (Johnston et al., 2021). Therefore, as a control for the possibility that injections of a control AAV with intrahippocampal EEG probe placement would be sufficient to lead to spontaneous seizures or cell death, PTEN/tdT mice received unilateral or bilateral injections of AAV-GFP into the dentate gyrus followed by continuous video EEG recordings (Table 5). A total of 4 mice were included in this cohort (Cohort 4). Two mice (33%) were excluded following anatomical analysis due to mistargeted viral transduction (Table 1). All survived each surgical procedure, and none of the mice with AAV-GFP injections developed spontaneous seizures (Fig. 6G, H, I).

**Figure 6.**
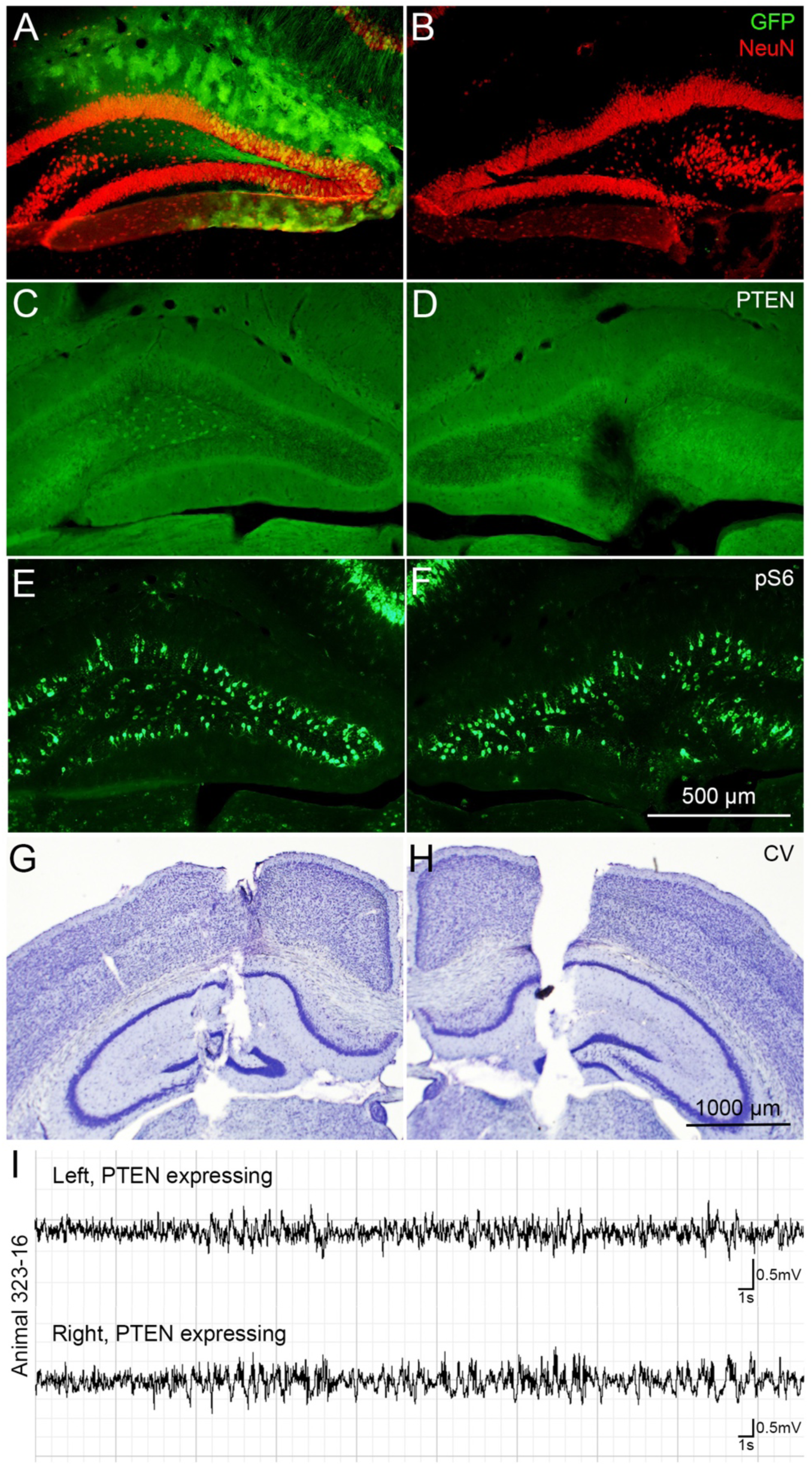
EEG recordings following AAV-GFP injection into the dentate gyrus of PTEN/tdT mice (Cohort 4). A) GFP expression in the ipsilateral (left) dentate gyrus of a PTEN/tdT mouse following unilateral AAV-GFP injection. B) Lack of GFP expression in the contralateral dentate gyrus. C-D) Preservation of PTEN expression in both hippocampi following unilateral AAV-GFP injection. E-F) pS6 immunoreactivity in the same control mouse. G-H) Cresyl violet stained sections show placement of bilateral electrodes in a control mouse. I) Representative EEG recordings from a mouse with AAV-GFP injection.

**Table 5:**
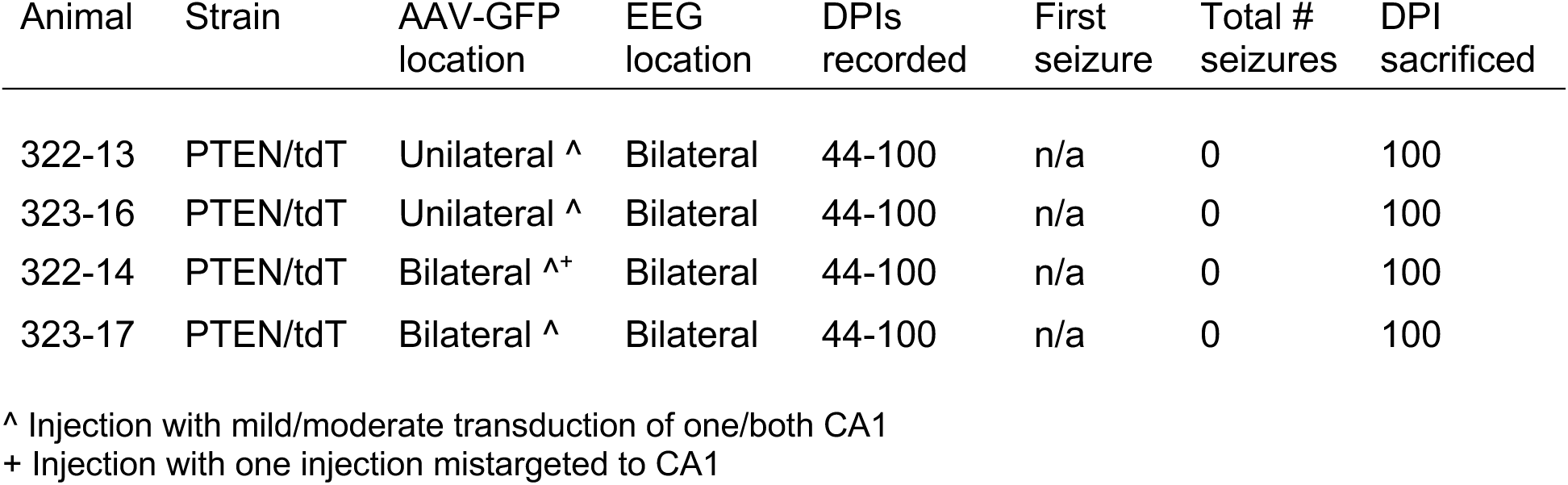
Cohort 4, Control mice used for EEG recordings.

Figure 6 illustrates representative images from a mouse with unilateral AAV-GFP injections recorded for 100 days post injection. In Figure 6A, AAV-driven GFP expression is evident throughout the dentate gyrus (green) and is lacking in the contralateral, non-injected dentate gyrus (Fig. 6B). Regardless of injection location, immunostaining for PTEN revealed intact PTEN expression in both hemispheres (Fig. 6C, D) and immunostaining for pS6 is comparable on the two sides of the brain (Fig. 6E, F) indicating no mTOR activation.

### 3.6 Extent of PTEN deletion correlates with seizure outcomes

Across Cohorts, PTEN deletion resulted in spontaneous seizures in 80, 67, and 100% of mice accessed (Fig. 7A), respectively. Despite our various approaches, there was no significant difference in the average latency to the first spontaneous seizure (Fig. 7B; one-way ANOVA: [F (2,10) = 2.489, p=0.1327]), the average total number of seizures (Fig. 7C; one-way ANOVA: [F (2,10) = 0.4214, p=0.6673]), or the average number of seizures per day (Fig. 7D, one-way ANOVA: [F (2,10) = 0.1313, p=0.8785]) across all cohorts. We therefore wondered if the amount of PTEN deletion in an individual animal would influence seizure outcomes. The mice with electrodes positioned contralateral to the PTEN deleted dentate gyrus in Cohort 3 allowed for detailed assessment of area of PTEN deletion without the complication of an implanted recording electrode. The area of transduction in these mice was characterized by a core in which most cells were tdT-positive, with transduction diminishing in the rostral and caudal directions (Fig. 8A). The average percent area transduced throughout the entire length of the dentate gyrus of PTEN/tdT mice was 36.16% +/− 6.79% (Fig. 8B).

**Figure 7.**
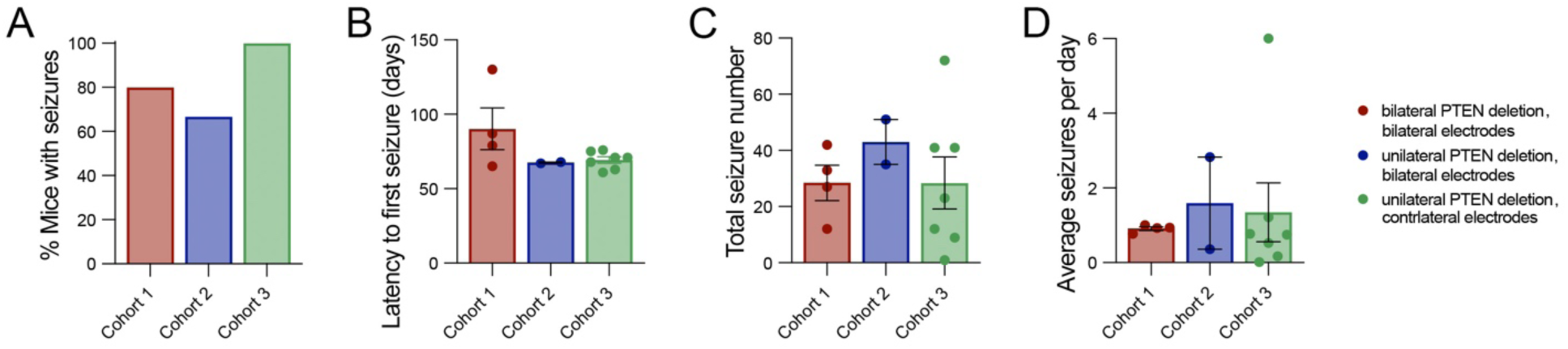
Seizure incidence across cohorts following PTEN deletion. A) Percent of mice within each cohort that displayed spontaneous seizures over the recording period. Mice with bilateral PTEN deletion and bilateral electrodes are shown in red (Cohort 1), mice with unilateral PTEN deletion and bilateral electrodes shown in blue (Cohort 2), and mice with unilateral PTEN deletion and contralateral electrode placement shown in green (Cohort 3). B) Latency to the first spontaneous seizure for each mouse within each cohort. C) Total seizure number across cohorts. D) Average number of seizures per day following PTEN deletion.

**Figure 8.**
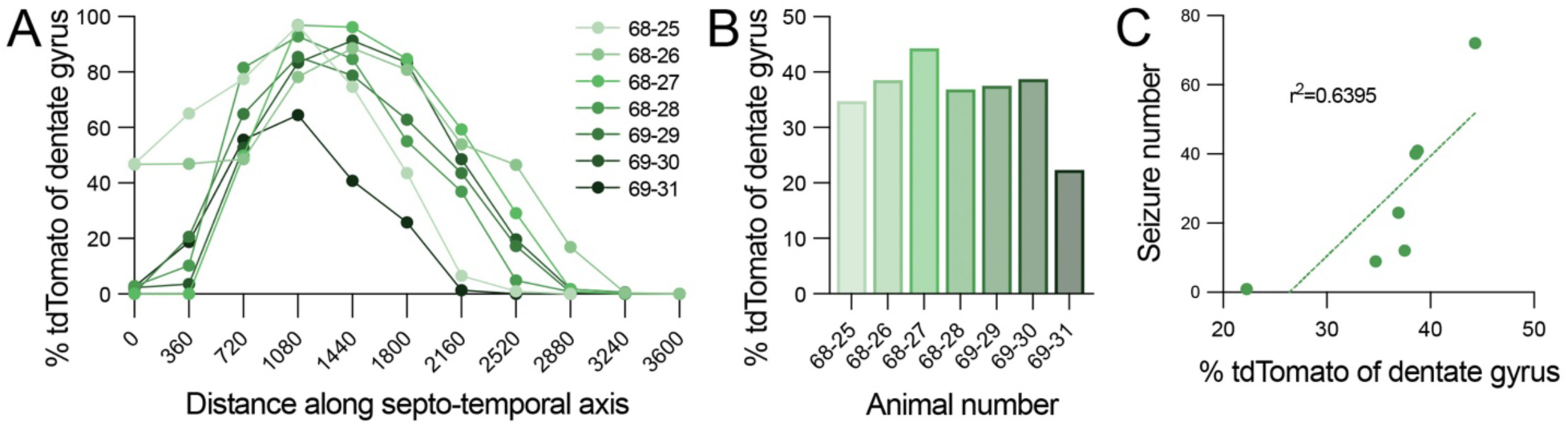
Relationship between PTEN deletion of the dentate gyrus and seizure number. A) Percent transduction of granule cells based on tdTomato expression by AAV-Cre over the septo-temporal axis of the ipsilateral dentate gyrus for each mouse in Cohort 3. B) Percent transduction of the entire ipsilateral dentate gyrus of the same mice. C) Relationship between percent transduction of the dentate gyrus for each mouse and cumulative number of seizures reported over the recording period, r^2^= 0.6395.

Interestingly, percent transduction was correlated with the total number of seizures in each mouse (r^2^=0.6395) where total number of seizures experienced over the recording period was greater in mice with larger areas of PTEN deletion within the dentate gyrus (Fig. 8C). Percent transduction was not correlated with the latency to the first seizure (r^2^=0.1728) or the average number of seizures per day (r^2^=0.06541) for each mouse (data not shown). In this cohort of mice, 2 of 7 mice had moderate transduction of CA1 pyramidal cells (Table 4); seizure number in these mice was within the range of the mice with transduction limited to the dentate gyrus, suggesting that some transduction of the CA1 did not alter seizure development.

### 3.7 Enlargement of granule cell bodies and processes after PTEN deletion

As in our other studies, a notable consequence of PTEN deletion was the enlargement of granule cell bodies and processes which was evident at 4 months post AAV-Cre injection. Figure 9A illustrates the enlargement of granule cell somata in the ipsilateral dentate gyrus (left) when compared to granule cells in the contralateral, PTEN-expressing dentate gyrus (right) in Cresyl violet stained sections. Soma cross sectional area was 127.50 +/− 15.93 μm^2^ for PTEN deleted granule cells vs. 67.96 +/− 14.19 μm^2^ for PTEN expressing granule cells (Fig. 9B). This approximately 2-fold increase in granule cell soma size is consistent to what we observed in our previous study (Yonan & Steward, 2023).

**Figure 9.**
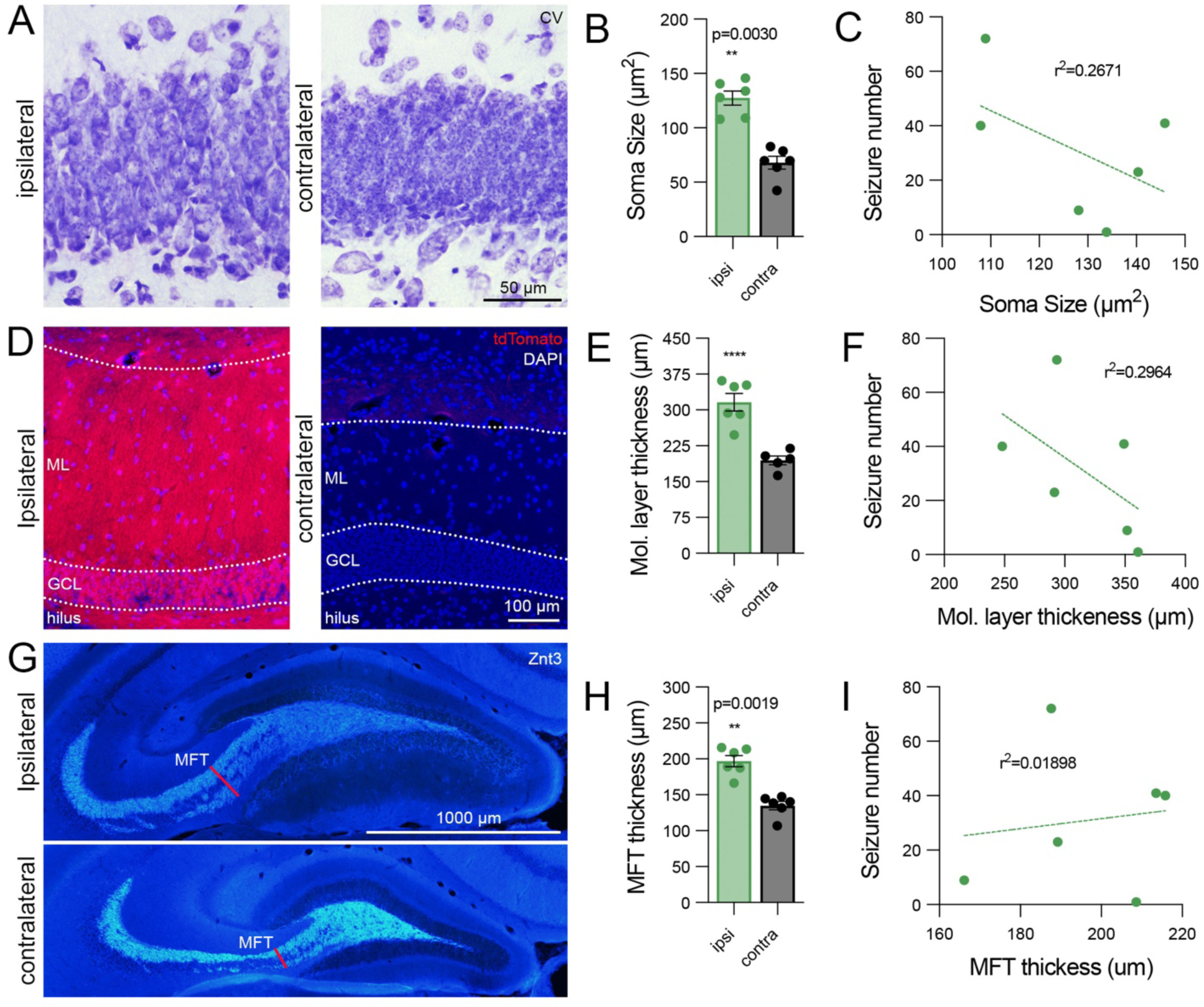
PTEN deletion triggers growth of granule cell bodies and processes. A) Representative Cresyl violet stained sections from the ipsilateral (left) and contralateral (right) dentate gyrus showing enlarged granule cell bodies. B) Average cell body size from the ipsilateral and contralateral dentate gyrus. Note each dot represents the average of 30 granule cells for an individual mouse. Sidak’s multiple comparisons for ipsilateral vs. contralateral sides in PTEN/tdT mice: p=0.0030. C) Relationship between soma size and seizure number for each animal, r^2^= 0.2671. D) Images of the molecular layer from the ipsilateral and contralateral dentate gyrus from the same mouse showing enlargement of the molecular layer, suggestive of increased apical dendrite length. E) Molecular layer thickness measured at the core of transduction for each mouse and the corresponding contralateral molecular layer. Sidak’s multiple comparisons for ipsilateral vs. contralateral sides in PTEN/tdT mice: p<0.0001. F) Relationship between molecular layer thickness and seizure number, r^2^= 0.2964. G) Znt3 labeling in the ipsilateral and contralateral dentate gyrus reveals enlargement of mossy fiber tract projections to the CA3. H) Measurements of mossy fiber tract thickness as it exits the hilus for each mouse (red lines labeled MFT in G). Sidak’s multiple comparisons for ipsilateral vs. contralateral sides in PTEN/tdT mice: p=0.0019. I) Relationship between mossy fiber tract thickness and seizure number, r^2^= 0.01898.

Enlargement of cell body size was accompanied by increases in the width of the molecular layer (Fig. 9D). We have previously confirmed that measurements of molecular layer thickness can serve as an indicator of apical dendrite length, given that dendrites typically extend to the hippocampal fissure (Yonan & Steward, 2023). Molecular layer thickness was 315.90 +/− 44.94 μm in the PTEN deleted dentate gyrus vs. 194.70 +/− 21.02 μm in the contralateral dentate gyrus (Fig. 9E), confirming PTEN-deletion induced expansion of dendritic arbors.

To assess whether PTEN deletion led to expansion of mossy fiber projections to CA3, sections at the core of transduction were stained for the zinc vesicular transporter, Znt3. As illustrated in Figure 9G, the collection of mossy fibers as they exit the hilus, which we term the mossy fiber tract (MFT) was notably thicker in the PTEN deleted dentate gyrus compared to the contralateral, PTEN-expressing dentate gyrus (MFT, red lines). Mossy fiber tract thickness was 196.80 +/− 19.30 μm in the ipsilateral dentate gyrus vs. 135.0 +/− 14.80 μm in the contralateral dentate gyrus (Fig. 9H). This expansion is again similar to what we observed in our previous study (Yonan & Steward, 2023).

Two-way ANOVA for all 3 parameters, collectively, revealed an overall significance for ipsilateral vs. contralateral sides [F (1,29) = 96.63, p<0.0001], a significance for measurement location [F (2,29) = 120.6, p<0.0001], and a significant interaction [F (2,29) = 5.816, p=0.0075]. Sidak’s multiple comparisons tests were also significant for somata, molecular layer, and mossy fiber tract measurements (see Figure 8 legend for statistics). No correlations were found between somata or process measurements and total seizure number for each animal (Fig. 9C, F, I).

### 3.8 Development of supra-granular mossy fibers following PTEN deletion

The growth of mossy fiber axons into the inner molecular layer, which indicates the formation of recurrent excitatory connections amongst granule cells, is a hallmark of different models of temporal lobe epilepsy (Althaus & Parent, 2012). We have previously reported the presence of supra-granular mossy fibers in some mice assessed at 4 months post PTEN deletion with greater than 40% transduction of the dentate gyrus (Yonan & Steward, 2023). We were therefore curious if similar ectopic projections would be seen in mice that display spontaneous seizures.

To quantify this, we used a modified scoring system from (Hunt et al., 2009) where a score of 0 indicates no mossy fibers in the granule cell layer and a score of 3 indicates dense mossy fiber staining in the inner molecular layer. Sections stained for Znt3 at the core of PTEN deletion for each mouse, as well as the contralateral dentate gyrus were assessed according to this metric.

At 4 months post deletion (Fig. 10A), 3 of 6 mice showed slight Znt3 labeling in the granule cell layer (score of 1), 2 of 6 mice displayed moderate Znt3 labeling in the granule cell layer with mild labeling in the inner molecular layer (score of 2), and 1 of 6 mice showed dense Znt3 labeling in the inner molecular layer (score of 3). Collectively 50% of mice (n=3) developed at least moderate Znt3 labeling in the inner molecular layer (score ≥ 2, Fig. 10B). All 3 mice had greater than 35% transduction of the dentate gyrus, although two mice with similar percent transductions only showed Znt3 labeling in the granule cell layer that did not extend into the inner molecular layer. Regression analysis of the relationship between percent transduction and mossy fiber score revealed a positive relationship (Fig. 10C, r^2^=0.4563). Of note, mice with higher mossy fiber scores experienced a greater total number of seizures over the recording period (Fig. 10D, r^2^=0.6244).

**Figure 10.**
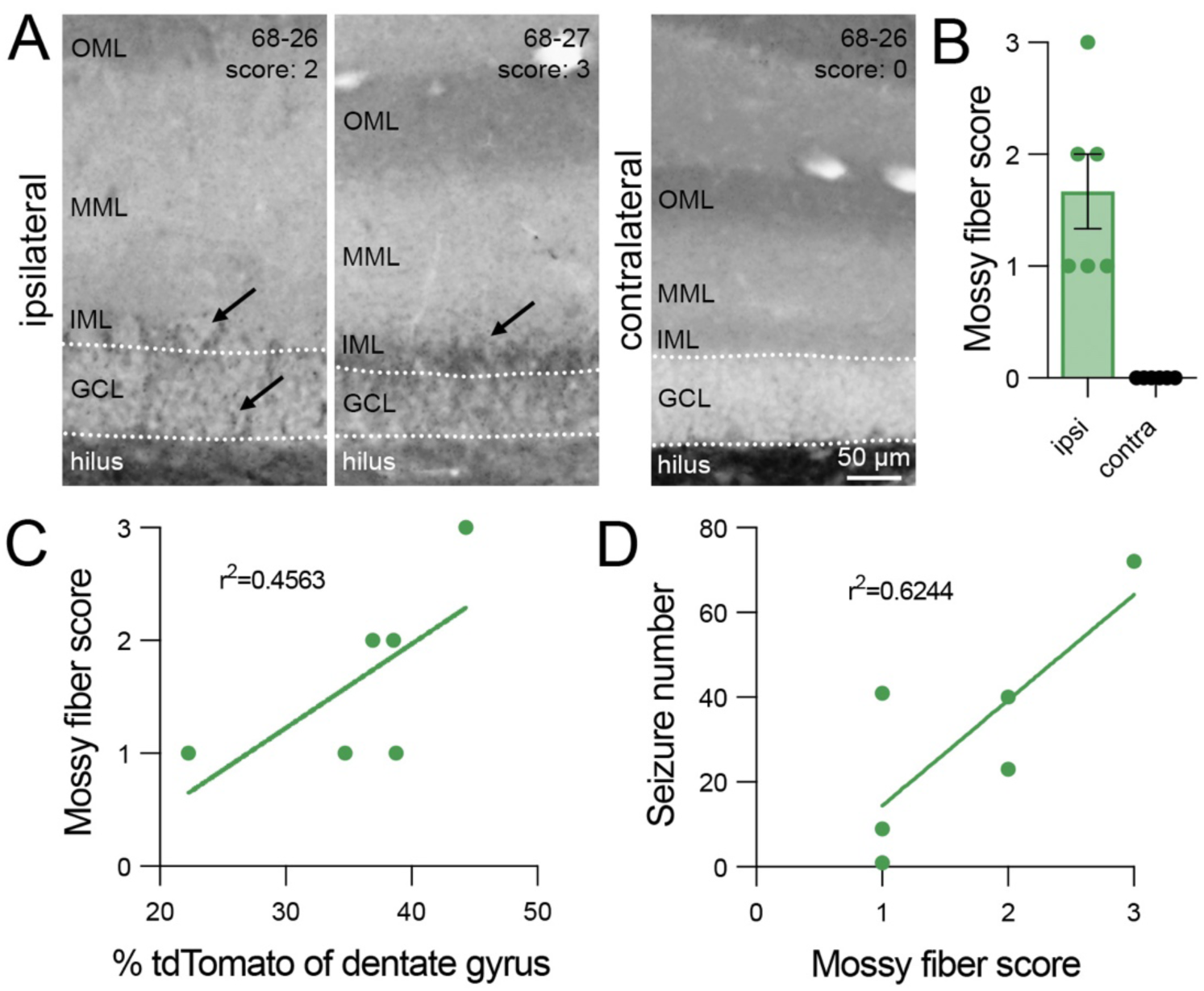
Presence of supragranular mossy fibers following PTEN deletion correlates with seizure number for each animal. A) Znt3 labeling in the granule cell layer and inner molecular layer reveals presence of supragranular mossy fibers following PTEN deletion (arrows) that are not present in the contralateral dentate gyrus. B) Mossy fiber score for the transduced and contralateral dentate gyrus of each mouse shows variability in the presence of supra-granular mossy fibers. Note representative scores are depicted in the top right corner of panel A. C) Relationship between percent transduction (PTEN deletion) and mossy fiber score, r^2^= 0.4563. D) Relationship between mossy fiber score and total seizure number, r^2^= 0.6244. Scoring scale: 0 - little to no Znt3 labeling in GCL, 1 - mild Znt3 labeling in GCL, 2 - moderate Znt3 labeling in GCL and mild labeling in IML, 3 - dense Znt3 labeling in IML.

### 3.9 Immunocytochemical evidence for seizures in mice without electrode implantation

The combined evidence above leaves little doubt that a focal area of PTEN deletion in the dentate gyrus of mature mice leads to the development of spontaneous seizures over time. The only small caveat is the implantation of a recording electrode, which could in theory act synergistically with PTEN deletion to increase excitability. As indirect evidence that seizures do occur following focal PTEN deletion, here we describe incidental findings of increased neuronal activation in a mouse without implanted recording electrodes that was part of an anatomical study in which tissue was collected 4-month post AAV-Cre injection. Of note, we did not monitor or record for behavioral seizures in this mouse.

We processed sections from this mouse as part of our assessment of s6 phosphorylation as an indicator of mTOR activation (above). In other mice, increases in the phosphorylation of ribosomal protein S6 in our model are confined to the region of PTEN deletion, and do not extend to regions of PTEN expression within the same dentate gyrus, or in the contralateral dentate gyrus (examples in Fig. 1E, F, Fig. 1M, N, Fig. 4E, F). In this mouse, however, there were striking increases in pS6 immunoreactivity in regions beyond the area of PTEN deletion in the ipsilateral dentate gyrus (Fig. 11A; PTEN vs. 11C, pS6) including in PTEN expressing granule cells in the contralateral dentate gyrus (Fig. 11B, D). Previous studies have reported prolonged neuronal activation after seizure activity in mouse models of temporal lobe epilepsy (Peng & Houser, 2005), and increased markers of mTOR signaling (Ahmed et al., 2021). Thus, a possible interpretation is that this mouse experienced a seizure in the period prior to tissue collection.

**Figure 11.**
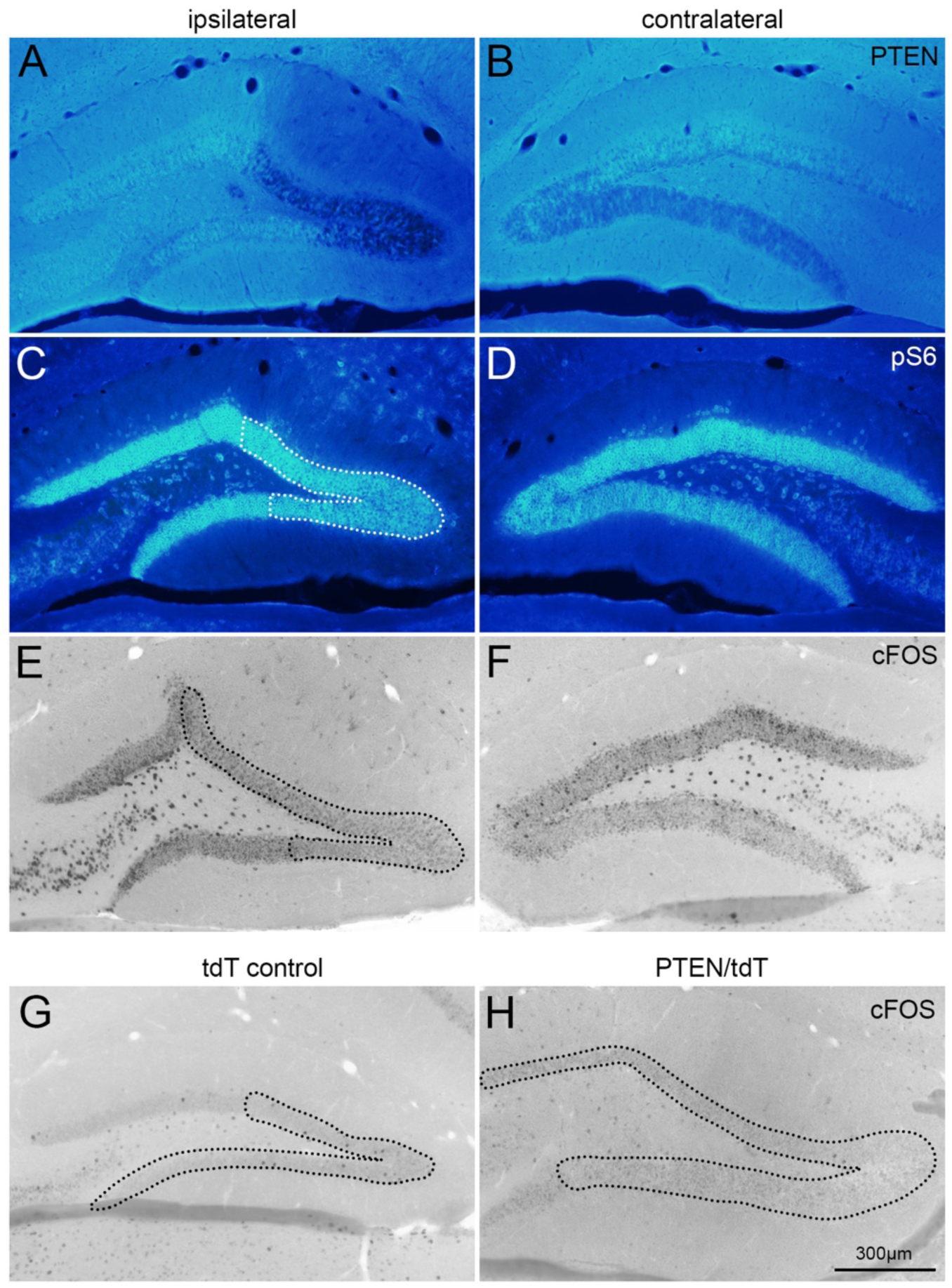
cFOS expression in the dentate gyrus of PTEN/tdT and control mice at 4 months following unilateral AAV-Cre injection. A) Effective PTEN deletion in the ipsilateral dentate gyrus of a PTEN/tdT mouse at 4 months after AAV-Cre injection. B) PTEN expression is maintained in the contralateral dentate gyrus. C) Increased phosphorylation of ribosomal protein S6 in the area of PTEN deletion (outlined in white) and in PTEN expression granule cells of the ipsilateral dentate gyrus. D) Increased phospho S6 in the contralateral dentate gyrus of the same mouse. E) Ipsilateral dentate gyrus of the same PTEN/tdT mouse showing increased cFos expression within and beyond the regions of PTEN deletion. F) Increased cFos expression in the contralateral dentate gyrus. G) Injections of AAV-Cre in a tdT control mouse does not result in increased cFOS expression (area of transduction outlined in black). H) PTEN deletion alone does not trigger increased cFOS expression (area of transduction and PTEN deletion outlined in black).

To further explore this possibility, we stained sections from this mouse for cFOS, which is strongly induced after a seizure; cFOS expression was also dramatically increased in both the ipsilateral (Fig. 11E) and contralateral dentate gyrus (Fig. 11F). In contrast, there are no increases in cFOS expression in dentate granule cells due to PTEN deletion alone (PTEN deleted area outline in Fig. 11H). Moreover, unilateral AAV-Cre injection into Rosa control mice does not trigger any notable increases in cFOS expression within the region of transduction (Fig. 11G). Thus, the dramatic increases in pS6 and cFOS immunoreactivity throughout the hippocampus suggest that this mouse may have experienced a seizure within hours before sacrifice.

### 3.10 Lack of hippocampal cell death following PTEN deletion

Epilepsy models that use convulsants to trigger spontaneous seizures following a latent period, like kainic acid and pilocarpine, often result in widespread neuronal death, especially in the CA3 region of the hippocampus (Curia et al., 2008; Drexel et al., 2012). We therefore wondered if seizures induced by adult, vector-mediated PTEN deletion would result in any notable hippocampal cell death. Figure 12 shows sample images from Cresyl violet-stained sections from PTEN/tdT mice injected with AAV-Cre (Fig. 12A) or AAV-GFP (Fig. 12B) near the center of viral transduction. There was no obvious cell loss in any hippocampal region with PTEN deletion (dentate gyrus in Fig. 12f), or in the neighboring CA1 (Fig. 12g) or CA3 (Fig. 12h) compared to the same hippocampal subregions in an AAV-GFP injected control (Fig. 12b-d). Importantly, the PTEN-deleted mouse shown here displayed the most seizures following unilateral PTEN deletion (Cohort 3, animal #68-27, 72 total seizures). There did appear to be fewer large neurons in the hilus in PTEN-deleted mice, which might indicate some loss or shrinkage of mossy cells within the hilus (compare arrowheads in Fig. 12f with Fig. 12b). Loss of mossy cells in the hilus often coincides with the presence of supra-granular mossy fibers and the formation of recurrent excitatory networks, both of which have been found to correlate with seizure frequency and duration (Hester & Danzer, 2013).

**Figure 12.**
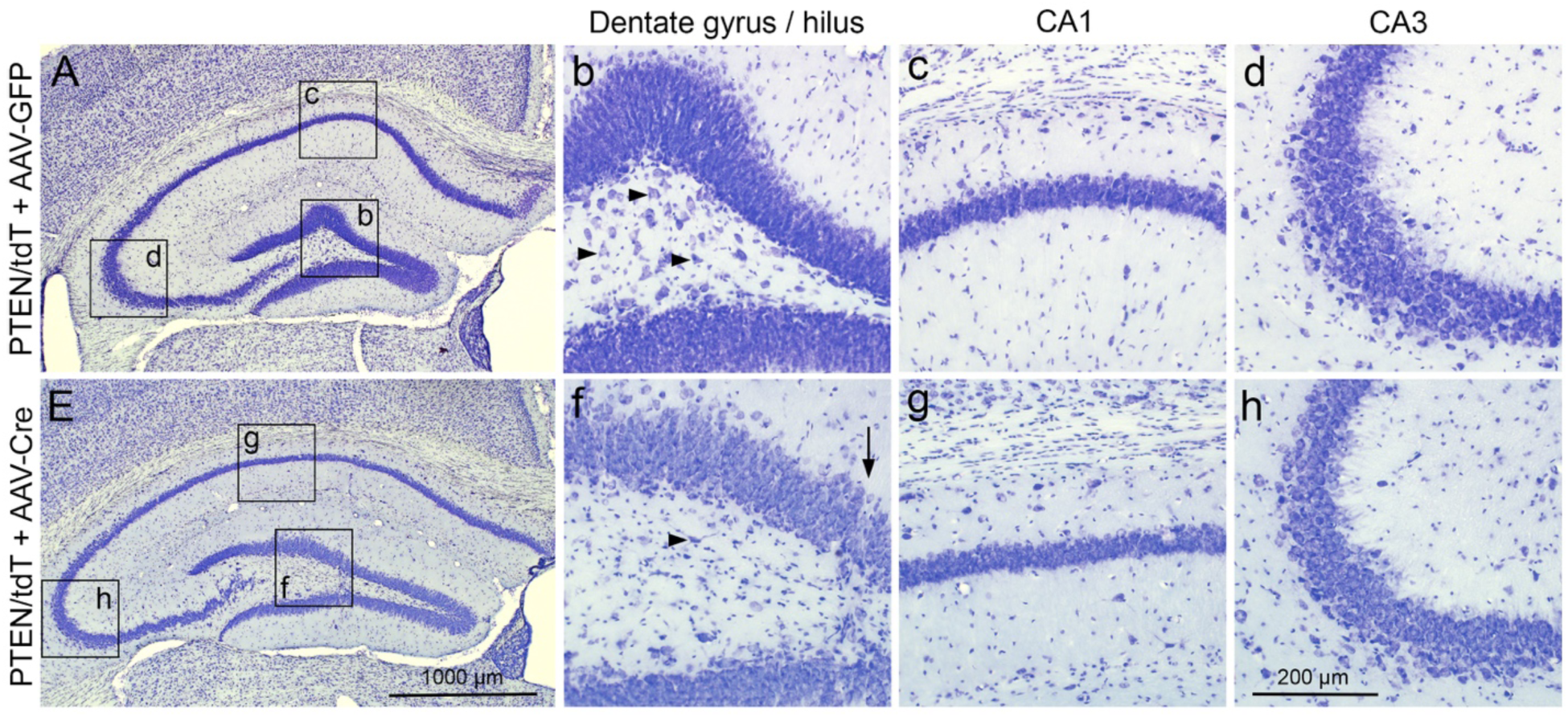
Lack of neuronal death in major hippocampal subregions following PTEN deletion. A) PTEN/tdT mouse injected with AAV-GFP into the dentate gyrus shows presence of neuronal cell bodies in the dentate gyrus and hilus (b), the CA1 (c), and the CA3 (d). B) Lack of neuronal cell death in the PTEN-deleted dentate gyrus (f), CA1 (g), and CA3 (h) following AAV-Cre injection in a PTEN/tdT mouse. Note fewer large neurons in the hilus in f (arrowheads vs. multiple arrowheads in b). Injection tract is observable in the PTEN deleted dentate gyrus in panel f.

### 3.11 Decreased immunostaining for GABA within the area of PTEN deletion

Interneurons located within the dentate gyrus provide feedforward and feedback inhibition onto granule cells. Loss of inhibition has been implicated in the development of seizures and reductions in the number of interneurons has been reported following developmental PTEN deletion (LaSarge et al., 2021). We therefore wondered if PTEN deletion in mature granule cells would result in decreases in GABA-ergic markers within the dentate gyrus. Immunostaining for glutamic acid decarboxylase, GAD67, a marker of GABA-expressing neurons and synapses, revealed subtle decreases in staining within the area of PTEN deletion in mice exhibiting spontaneous seizures (Fig. 13A) in comparison to the contralateral, PTEN expressing dentate gyrus (Fig. 13B). These decreases in staining could reflect an actual slight decrease in GABAergic innervation or could be due to the fact that immunostaining is diluted as a consequence of the overall enlargement of the dentate gyrus (essentially an increase in the denominator of density/unit area).

**Figure 13.**
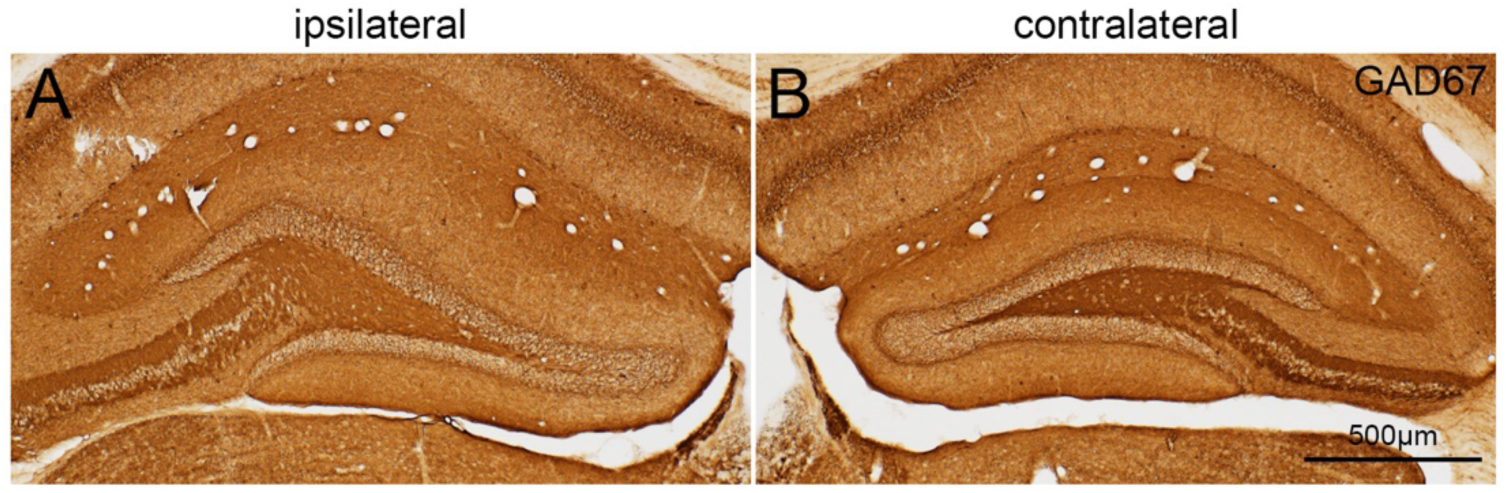
GAD67 immunoreactivity following vector-mediated PTEN deletion. A) Decreased GAD67 immunoreactivity in the granule cell layer, molecular layer, and hilus in the area of PTEN deletion. B) Control GAD67 labeling in the contralateral dentate gyrus.

## 4.0 Discussion

The goal of this study was to explore whether PTEN deletion and resulting growth of granule cells and alterations in hippocampal circuitry result in the development of spontaneous seizures. The principal findings are: (1) deletion of PTEN in adult dentate gyrus granule cells results in delayed development of recurrent spontaneous seizures (epilepsy) in the majority of mice; seizures were not observed in mice that received AAV-GFP (controls); (2) the latency to the first spontaneous seizure is about two months, (3) spontaneous seizures develop with either a bilateral or unilateral focus of deletion; (4) in contrast to excitotoxin models, epileptogenesis is not associated with obvious neuron loss in CA3. Together these results document a novel, convulsant/toxin-free model of hippocampal/temporal lobe epileptogenesis.

### 4.1 Focal, unilateral PTEN deletion is sufficient for the development of spontaneous, recurrent hippocampus-origin seizures

Continuous video-EEG monitoring with bilateral, intrahippocampal electrodes revealed that either bilateral or unilateral PTEN deletion in the dentate gyrus led to the development of spontaneous electrographic and behavioral seizures in 80% and 67% of mice, respectively. Seizures originated in either hippocampus in mice with bilateral PTEN deletion, but consistently originated in the PTEN deleted dentate gyrus with unilateral PTEN deletion.

All mice that received unilateral injections of AAV-Cre into the dentate gyrus with recording electrodes positioned in the contralateral hippocampus also developed spontaneous seizures. This largely excludes the possibility that seizure development was due to the combination of PTEN deletion and electrode implantation. Such a combinatorial effect has been suggested with electrode implantation in kainic acid and pilocarpine models of epilepsy (Balzekas et al., 2016; Levesque et al., 2016).

Importantly, spontaneous seizures developed with as little as 22% transduction of the dentate gyrus on one side, indicating that a relatively small, unilateral focus of PTEN deletion is sufficient for bilateral electrographic and behavioral seizures. This is reminiscent of the situation in human TLE where a unilateral focus can eventually lead to generalized seizures.

Transduction and PTEN deletion was not restricted to the dentate gyrus in some mice, which could contribute to seizure development. However, 6 mice with well targeted unilateral PTEN deletion developed seizures (1 with bilateral electrodes and 5 with contralateral electrode placement). In addition, seizures were neither more frequent nor more severe in mice with transduction involving the CA1 region in addition to the dentate gyrus.

### 4.2 Latent period for seizure development

Spontaneous seizures were not observed in mice until about 2 months following PTEN deletion, regardless of pattern of PTEN deletion or electrode placement. This delayed onset of spontaneous seizures, or latent period, suggests progressive network modification over time. Of note, epileptogenesis due to focal PTEN deletion follows a different time course than the commonly used kainic acid and pilocarpine models of TLE, which have relatively short latent periods, ranging from 10-30 and 4-40 days, respectively, before the appearance of spontaneous seizures (see review by (Levesque et al., 2016)). The delayed development of seizures with PTEN deletion is reminiscent of the delayed development of temporal lobe epilepsy (TLE) after some insult or injury involving the hippocampus in humans which can last several years (Buckmaster, 2004).

### 4.3 Relationship between seizure development and growth of granule cells

PTEN deletion triggers two time-dependent processes; 1) growth of dentate granule cell bodies, elongation of dendrites and new spine formation, and expansion of axonal projections; 2) development of seizures. In our previous study, increases in granule cell body size were statistically significant at 2 months (when seizures appear) but granule cell size continued to increase over time. Dendritic and axonal elongation and alterations in connectivity were not evident until 4 months post-deletion (Yonan & Steward, 2023). Because seizures develop before growth responses are fully developed, seizures may be due to alterations in neuron-intrinsic physiological processes due to PTEN deletion. Alternatively, it is possible that early morphological changes are sufficient to trigger an epileptogenic circuit. Further studies will be required to explore these possibilities.

### 4.4 Sudden death during seizures

Of the 13 mice that developed seizures, 3 eventually died during a seizure (23%). This is in contrast to reports of high mortality rates that result from increasing seizure severity with PTEN deletion in early development (Kwon et al., 2003; Kwon et al., 2001; Matsushita et al., 2016; Pun et al., 2012; Sunnen et al., 2011). One possible explanation is that seizures in our model are less severe, and thus less likely to lead to respiratory arrest. Other possibilities include some compensatory mechanisms or network properties in mature circuits that attenuate seizure progression.

Of note, mortality rates in the present study are somewhat higher than in our previous study with the same PTEN deletion model but without electrode implantation (Yonan & Steward, 2023). It is possible that seizures are more severe and/or that mice are more susceptible to respiratory arrest when PTEN deletion and electrode implantation are combined.

### 4.5 Supra-granular mossy fibers not required, but may contribute to seizure outcomes

Supragranular mossy fibers develop in excitotoxin models of epilepsy and are thought to represent the formation of a recurrent excitatory circuit. Supragranular mossy fibers were noted in studies involving PTEN deletion in early development (Amiri et al., 2012; Kwon et al., 2006; Pun et al., 2012; Sunnen et al., 2011), and were also seen in 40% of mice at 4 and 6 months after PTEN deletion in our previous study (Yonan & Steward, 2023). In the present study, all mice with unilateral PTEN deletion with contralateral electrode placement developed seizures and showed some mossy fiber labeling in the granule cell layer, but only 50% of mice had actual supragranular mossy fibers. Thus, spontaneous seizures can occur in the absence of supragranular mossy fibers as reported in studies of developmental PTEN deletion (Pun et al., 2012), but may contribute to seizure progression over time.

### 4.6 New model of epileptogenesis and adult-onset epilepsy

Various laboratory models have been developed to investigate mechanisms of epileptogenesis, including brain injury, hypoxia and ischemia, kindling, and chemical convulsants [for review see (Buckmaster, 2004)]. Many models including kainic acid and pilocarpine models trigger extensive neuronal death both within and beyond the hippocampus (Curia et al., 2008; Drexel et al., 2012). Whether neuronal death is required for epileptogenesis during development and in adulthood is up for debate (Baram et al., 2011). For example, febrile status epilepticus (FSE) often results in epilepsy without widespread cell loss (Dube et al., 2010). Our model of vector-mediated focal PTEN deletion in the adult dentate gyrus provides another model of epileptogenesis without apparent neuronal death, which may provide a unique opportunity to define different mechanisms of adult-onset temporal lobe epilepsy than have not been explored in other animal models.

## Acknowledgements

We would like to thank Dr. Alicia Hall, Karla McHale and Tram Ngyuen for their technical support and contributions to this study. This work was supported by NIH-NS108189 (OS), NS108296 (TZB, KDC). J. Yonan was supported by NIH T32 Epilepsy training grant NS045540 (TZB, PI), NIH R01 Diversity Supplement to NIH-NS108189, a dissertation year fellowship from the University of California, Irvine, School of Medicine and the University of California Chancellor’s Postdoctoral Fellowship.

## REFERENCES

Ahmed, M. M., Carrel, A. J., Cruz Del Angel, Y., Carlsen, J., Thomas, A. X., Gonzalez, M. I., Gardiner, K. J., & Brooks-Kayal, A. (2021). Altered Protein Profiles During Epileptogenesis in the Pilocarpine Mouse Model of Temporal Lobe Epilepsy. Front Neurol, 12, 654606. 10.3389/fneur.2021.654606

Althaus, A. L., & Parent, J. M. (2012, Sep 20). Pten-less dentate granule cells make fits. Neuron, 75(6), 938–940. 10.1016/j.neuron.2012.09.008

Amiri, A., Cho, W., Zhou, J., Birnbaum, S. G., Sinton, C. M., McKay, R. M., & Parada, L. F. (2012, Apr 25). Pten deletion in adult hippocampal neural stem/progenitor cells causes cellular abnormalities and alters neurogenesis. J Neurosci, 32(17), 5880–5890. 10.1523/JNEUROSCI.5462-11.2012

Arafa, S. R., LaSarge, C. L., Pun, R. Y. K., Khademi, S., & Danzer, S. C. (2019, Jan). Self-reinforcing effects of mTOR hyperactive neurons on dendritic growth. Exp Neurol, 311, 125–134. 10.1016/j.expneurol.2018.09.019

Backman, S. A., Stambolic, V., Suzuki, A., Haight, J., Elia, A., Pretorius, J., Tsao, M. S., Shannon, P., Bolon, B., Ivy, G. O., & Mak, T. W. (2001, Dec). Deletion of Pten in mouse brain causes seizures, ataxia and defects in soma size resembling Lhermitte-Duclos disease. Nat Genet, 29(4), 396–403. 10.1038/ng782

Balzekas, I., Hernandez, J., White, J., & Koh, S. (2016, May 27). Confounding effect of EEG implantation surgery: Inadequacy of surgical control in a two hit model of temporal lobe epilepsy. Neurosci Lett, 622, 30–36. 10.1016/j.neulet.2016.04.033

Baram, T. Z., Jensen, F. E., & Brooks-Kayal, A. (2011, Jan). Does acquired epileptogenesis in the immature brain require neuronal death. Epilepsy Curr, 11(1), 21–26. 10.5698/1535-7511-11.1.21

Buckmaster, P. S. (2004, Oct). Laboratory animal models of temporal lobe epilepsy. Comp Med, 54(5), 473–485. https://www.ncbi.nlm.nih.gov/pubmed/15575361

Chen, K. D., Hall, A. M., Garcia-Curran, M. M., Sanchez, G. A., Daglian, J., Luo, R., & Baram, T. Z. (2021, Mar). Augmented seizure susceptibility and hippocampal epileptogenesis in a translational mouse model of febrile status epilepticus. Epilepsia, 62(3), 647–658. 10.1111/epi.16814

Curia, G., Longo, D., Biagini, G., Jones, R. S., & Avoli, M. (2008, Jul 30). The pilocarpine model of temporal lobe epilepsy. J Neurosci Methods, 172(2), 143–157. 10.1016/j.jneumeth.2008.04.019

Drexel, M., Preidt, A. P., & Sperk, G. (2012, Oct). Sequel of spontaneous seizures after kainic acid-induced status epilepticus and associated neuropathological changes in the subiculum and entorhinal cortex. Neuropharmacology, 63(5), 806–817. 10.1016/j.neuropharm.2012.06.009

Dube, C., Richichi, C., Bender, R. A., Chung, G., Litt, B., & Baram, T. Z. (2006, Apr). Temporal lobe epilepsy after experimental prolonged febrile seizures: prospective analysis. Brain, 129(Pt 4), 911–922. 10.1093/brain/awl018

Dube, C. M., Ravizza, T., Hamamura, M., Zha, Q., Keebaugh, A., Fok, K., Andres, A. L., Nalcioglu, O., Obenaus, A., Vezzani, A., & Baram, T. Z. (2010, Jun 2). Epileptogenesis provoked by prolonged experimental febrile seizures: mechanisms and biomarkers. J Neurosci, 30(22), 7484–7494. 10.1523/JNEUROSCI.0551-10.2010

Fraser, M. M., Bayazitov, I. T., Zakharenko, S. S., & Baker, S. J. (2008, Jan 24). Phosphatase and tensin homolog, deleted on chromosome 10 deficiency in brain causes defects in synaptic structure, transmission and plasticity, and myelination abnormalities. Neuroscience, 151(2), 476–488. 10.1016/j.neuroscience.2007.10.048

Fraser, M. M., Zhu, X., Kwon, C. H., Uhlmann, E. J., Gutmann, D. H., & Baker, S. J. (2004, Nov 1). Pten loss causes hypertrophy and increased proliferation of astrocytes in vivo. Cancer Res, 64(21), 7773–7779. 10.1158/0008-5472.CAN-04-2487

Gallent, E. A., & Steward, O. (2018, May). Neuronal PTEN deletion in adult cortical neurons triggers progressive growth of cell bodies, dendrites, and axons. Exp Neurol, 303, 12–28. 10.1016/j.expneurol.2018.01.005

Garcia-Curran, M. M., Hall, A. M., Patterson, K. P., Shao, M., Eltom, N., Chen, K., Dube, C. M., & Baram, T. Z. (2019, Nov/Dec). Dexamethasone Attenuates Hyperexcitability Provoked by Experimental Febrile Status Epilepticus. eNeuro, 6(6). 10.1523/ENEURO.0430-19.2019

Hester, M. S., & Danzer, S. C. (2013, May 22). Accumulation of abnormal adult-generated hippocampal granule cells predicts seizure frequency and severity. J Neurosci, 33(21), 8926–8936. 10.1523/JNEUROSCI.5161-12.2013

Hunt, R. F., Scheff, S. W., & Smith, B. N. (2009, Feb). Posttraumatic epilepsy after controlled cortical impact injury in mice. Exp Neurol, 215(2), 243–252. 10.1016/j.expneurol.2008.10.005

Johnston, S., Parylak, S. L., Kim, S., Mac, N., Lim, C., Gallina, I., Bloyd, C., Newberry, A., Saavedra, C. D., Novak, O., Goncalves, J. T., Gage, F. H., & Shtrahman, M. (2021, Jul 14). AAV ablates neurogenesis in the adult murine hippocampus. Elife, 10. 10.7554/eLife.59291

Kwon, C. H., Luikart, B. W., Powell, C. M., Zhou, J., Matheny, S. A., Zhang, W., Li, Y., Baker, S. J., & Parada, L. F. (2006, May 4). Pten regulates neuronal arborization and social interaction in mice. Neuron, 50(3), 377–388. 10.1016/j.neuron.2006.03.023

Kwon, C. H., Zhu, X., Zhang, J., & Baker, S. J. (2003, Oct 28). mTor is required for hypertrophy of Pten-deficient neuronal soma in vivo. Proc Natl Acad Sci U S A, 100(22), 12923–12928. 10.1073/pnas.2132711100

Kwon, C. H., Zhu, X., Zhang, J., Knoop, L. L., Tharp, R., Smeyne, R. J., Eberhart, C. G., Burger, P. C., & Baker, S. J. (2001, Dec). Pten regulates neuronal soma size: a mouse model of Lhermitte-Duclos disease. Nat Genet, 29(4), 404–411. 10.1038/ng781

LaSarge, C. L., Pun, R. Y., Muntifering, M. B., & Danzer, S. C. (2016, Dec). Disrupted hippocampal network physiology following PTEN deletion from newborn dentate granule cells. Neurobiol Dis, 96, 105–114. 10.1016/j.nbd.2016.09.004

LaSarge, C. L., Pun, R. Y. K., Gu, Z., Riccetti, M. R., Namboodiri, D. V., Tiwari, D., Gross, C., & Danzer, S. C. (2021, May). mTOR-driven neural circuit changes initiate an epileptogenic cascade. Prog Neurobiol, 200, 101974. 10.1016/j.pneurobio.2020.101974

LaSarge, C. L., Santos, V. R., & Danzer, S. C. (2015, Mar). PTEN deletion from adult-generated dentate granule cells disrupts granule cell mossy fiber axon structure. Neurobiol Dis, 75, 142–150. 10.1016/j.nbd.2014.12.029

Levesque, M., Avoli, M., & Bernard, C. (2016, Feb 15). Animal models of temporal lobe epilepsy following systemic chemoconvulsant administration. J Neurosci Methods, 260, 45–52. 10.1016/j.jneumeth.2015.03.009

Matsushita, Y., Sakai, Y., Shimmura, M., Shigeto, H., Nishio, M., Akamine, S., Sanefuji, M., Ishizaki, Y., Torisu, H., Nakabeppu, Y., Suzuki, A., Takada, H., & Hara, T. (2016, Mar 10). Hyperactive mTOR signals in the proopiomelanocortin-expressing hippocampal neurons cause age-dependent epilepsy and premature death in mice. Sci Rep, 6, 22991. 10.1038/srep22991

Peng, Z., & Houser, C. R. (2005, Aug 3). Temporal patterns of fos expression in the dentate gyrus after spontaneous seizures in a mouse model of temporal lobe epilepsy. J Neurosci, 25(31), 7210–7220. 10.1523/JNEUROSCI.0838-05.2005

Pitkanen, A., Nissinen, J., Nairismagi, J., Lukasiuk, K., Grohn, O. H., Miettinen, R., & Kauppinen, R. (2002). Progression of neuronal damage after status epilepticus and during spontaneous seizures in a rat model of temporal lobe epilepsy. Prog Brain Res, 135, 67–83. 10.1016/S0079-6123(02)35008-8

Pun, R. Y., Rolle, I. J., Lasarge, C. L., Hosford, B. E., Rosen, J. M., Uhl, J. D., Schmeltzer, S. N., Faulkner, C., Bronson, S. L., Murphy, B. L., Richards, D. A., Holland, K. D., & Danzer, S. C. (2012, Sep 20). Excessive activation of mTOR in postnatally generated granule cells is sufficient to cause epilepsy. Neuron, 75(6), 1022–1034. 10.1016/j.neuron.2012.08.002

Santos, V. R., Pun, R. Y. K., Arafa, S. R., LaSarge, C. L., Rowley, S., Khademi, S., Bouley, T., Holland, K. D., Garcia-Cairasco, N., & Danzer, S. C. (2017, Dec). PTEN deletion increases hippocampal granule cell excitability in male and female mice. Neurobiol Dis, 108, 339–351. 10.1016/j.nbd.2017.08.014

Steward, O., Coulibay, A., Metcalfe, M., Yonan, J. M., & Yee, K. M. (2019). AAVshRNA-mediated PTEN knockdown in adult neurons attenuates activity-dependent immediate early gene induction. Exp Neurol. 10.1016/j.expneurol.2019.113098

Sunnen, C. N., Brewster, A. L., Lugo, J. N., Vanegas, F., Turcios, E., Mukhi, S., Parghi, D., D’Arcangelo, G., & Anderson, A. E. (2011, Nov). Inhibition of the mammalian target of rapamycin blocks epilepsy progression in NS-Pten conditional knockout mice. Epilepsia, 52(11), 2065–2075. 10.1111/j.1528-1167.2011.03280.x

Williams, M. R., DeSpenza, T., Jr., Li, M., Gulledge, A. T., & Luikart, B. W. (2015, Jan 21). Hyperactivity of newborn Pten knock-out neurons results from increased excitatory synaptic drive. J Neurosci, 35(3), 943–959. 10.1523/JNEUROSCI.3144-14.2015

Yonan, J. M., & Steward, O. (2023, Aug). Vector-mediated PTEN deletion in the adult dentate gyrus initiates new growth of granule cell bodies and dendrites and expansion of mossy fiber terminal fields that continues for months. Neurobiol Dis, 184, 106190. 10.1016/j.nbd.2023.106190

